# Illusory sound texture reveals multi-second statistical completion in auditory scene analysis

**DOI:** 10.1101/681965

**Authors:** Richard McWalter, Josh H. McDermott

**Affiliations:** Department of Brain and Cognitive Sciences, MIT; Center for Brains, Minds and Machines, MIT; McGovern Institute for Brain Research, MIT; Program in Speech and Hearing Biosciences and Technology, Harvard University

## Abstract

Sound sources in the world are experienced as stable even when intermittently obscured, implying perceptual completion mechanisms that “fill in” missing sensory information. We demonstrate a filling-in phenomenon in which the brain extrapolates the statistics of background sounds (textures) over periods of several seconds when they are interrupted by another sound, producing vivid percepts of illusory texture. The effect differs from previously described completion effects in that 1) the extrapolated sound must be defined statistically given the stochastic nature of texture, and 2) in lasting much longer, enabling introspection and facilitating assessment of the underlying representation. Illusory texture appeared to be integrated into texture statistic estimates indistinguishably from actual texture, suggesting that it is represented similarly to actual texture. The illusion appears to represent an inference about whether the background is likely to continue during concurrent sounds, providing a stable representation of the environment despite unstable sensory evidence.

## Introduction

Perception consists of inferences about the state of the world given sensory stimulation. These inferences are typically ill-posed, akin to solving an equation with multiple unknowns, as when estimating sound sources from a mixture of sources, or three-dimensional depth relationships from two-dimensional images. As such, most perceptual inferences necessitate assumptions about the world variables being estimated – knowledge of environmental regularities that have been internalized over development or evolution. Such inferences and their constraining assumptions can be revealed by illusions – artificial situations or stimuli that are erroneously perceived, illustrating the assumptions that produce correct inferences most of the time in natural environments.

One class of perceptual inferences, and associated illusions, occur when perceptual systems “fill in” missing data, as when objects are occluded or sound sources are masked. Sensory traces of objects and sounds are physically interrupted in such situations, and yet we usually experience them as continuous. Classic examples in vision include illusory contours^1^, often termed “modal” completion because the contours are subjectively visible, and “amodal” completion^2^, when an object is seen to continue beneath an occluding surface (Figure 1a). In audition, perceptual completion is known to occur for speech and tone-like sounds^3–12^, which are heard as continuous even when brief segments are removed and replaced with noise, provided that the noise is sufficiently intense as to plausibly mask the speech or tone (Figure 1b). Because the stimuli in such “continuity illusions” are equally consistent with discontinuous sound sources or objects, they appear to represent an inference that the sound source was likely to have continued during the noise. However, previously documented effects are for the most part short lasting, spanning brief gaps of a few hundred milliseconds, and have mostly been limited to sounds that might be produced by individual sound sources in the world.

**Figure 1.**
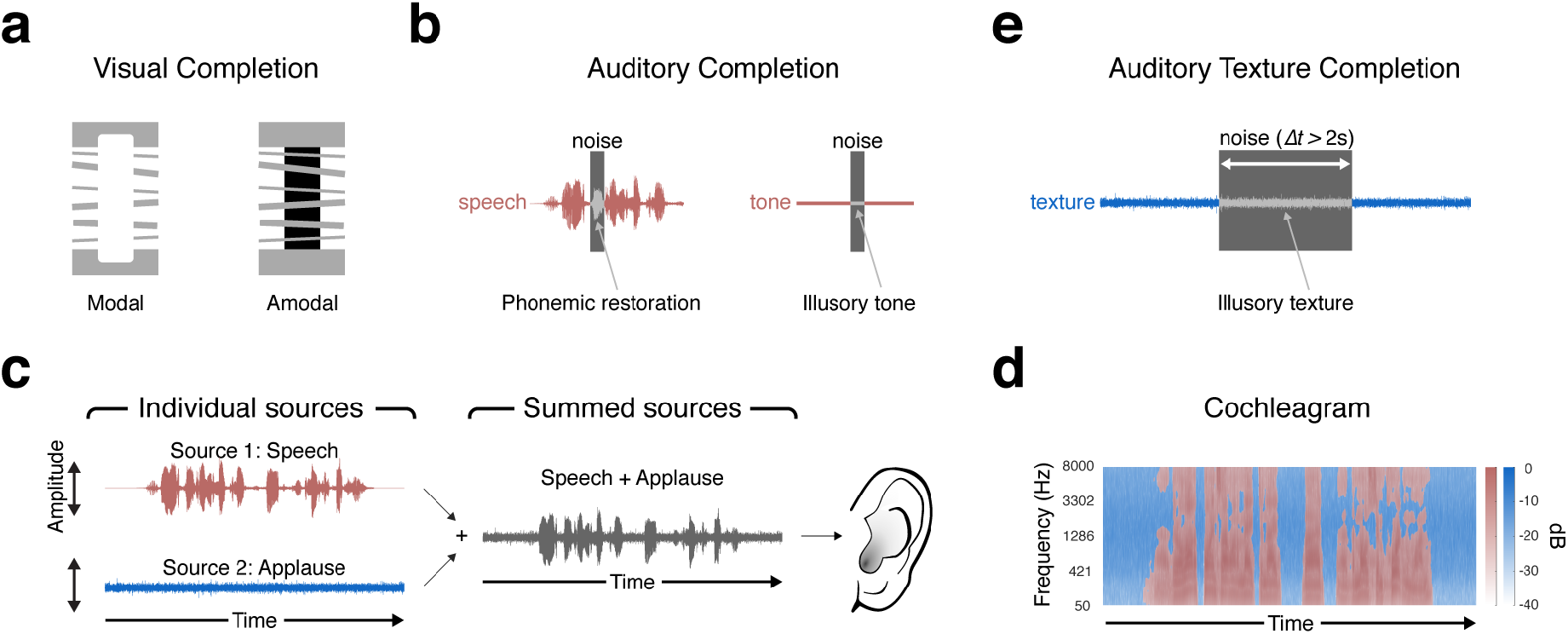
Visual/Auditory Completion and Auditory Scenes. **(a)** Examples of “modal” and “amodal” visual completion. In the modal case (left), we see diagonal white bars overlaid on the cube. However, the bar edges do not produce contrast where they overlap the background. In the amodal case (right), the grey bars are seen to continue behind the cube. **(b)** Traditional examples of auditory completion. Tones can be heard to continue when interrupted with a sound (e.g. noise) sufficiently loud as to plausibly mask the tone. The tone is heard to continue even though it is not physically present during the noise. Illusory phonemes (or phonemic restoration) can also occur under similar conditions, when a short segment of speech is replaced with a louder sound. Here and in e, the grey shading of the sound waveform symbolizes illusory sound that is heard despite not being present in the stimulus. **(c)** Example auditory scene in which two sound sources (speech and applause) combine to form a single waveform that arrives at the ear. **(d)** Masking of background sound texture. The speech sound has more energy during certain time-frequency windows, intermittently masking the background applause sound. The red areas show windows where the speech has more energy than the applause. The blue areas show windows where the applause has more sound energy than the speech. **(e)** Illusory texture from auditory texture completion. Illusory texture can be heard when a texture is interrupted with another sound sufficiently loud as to mask the texture. The texture is heard to continue even though it is not physically present during the interrupting sound. Texture completion is differentiated from other forms of auditory completion in the long temporal extent over which illusory textures can be heard (often exceeding 2 seconds).

Auditory scenes frequently contain sound textures – sounds produced by the superposition of many sound sources, such as a room of people talking or clapping, swarms of insects, or rain^13,14^. Sound textures often continue over long periods of time, providing a background to an auditory scene. Given that the background is often physically continuous but intermittently obscured by other sound sources (Figure 1c&d), we investigated whether textures might be inferred to continue during such interruptions. Texture completion seemed potentially interesting in part because textures are believed to be represented with statistics that summarize acoustic information over time^14–21^, raising the question of whether statistical representations could be “filled in”.

Here we describe a class of illusions in which texture subjectively continues when followed by sufficiently loud masking noise, producing vivid illusory percepts (Figure 1e). Examples of the illusion can be found here: http://mcdermottlab.mit.edu/textcont.html

Unlike the brief illusory continuity heard for speech, tones or other non-stationary sounds, illusory textures persisted for up to several seconds. Because textures are dense, stochastic, and in many cases defined only by summary statistics, the perceptual completion appears to be mediated by extrapolated statistics. The extended duration of the illusory sound facilitated investigation of the underlying representation. Illusory texture appears to be incorporated into the statistical estimation process for texture, biasing the perception of subsequent textures in the same manner as physically realized texture. This effect suggests that the underlying perceptual completion processes instantiate representations like those produced by actual texture. The results reveal a form of perceptual completion that is statistical in nature, extends over periods of seconds, and that appears to be specific to dense stationary sounds, suggesting a difference in the representation of textures and discrete events.

## Results

We conducted a series of experiments to document the illusion, establish the conditions in which it occurs, and probe its representational basis. We first set out to validate our informal observations of illusory texture. We asked listeners to judge the continuity of a variety of sounds interrupted by noise, exploring whether the effect was specific to textures (Experiment 1). We then measured the temporal extent of the illusion (Experiments 2 and 3). Next, we characterized the relationship of the illusion to masking (Experiment 4) and physical continuity (Experiment 5). We used the temporal integration of texture to ask if illusory texture is represented like actual texture (Experiment 6). Finally, we explored whether illusory texture could also occur when textures were part of an auditory scene with other sounds (Experiment 7).

### Experiment 1: Illusory texture occurs for most stationary sounds

As an initial validation of our subjective observations, we asked listeners to judge the continuity of a variety sounds interrupted by noise. We selected a set of 80 ‘inducer’ sounds that included some textures as well as a wide assortment of non-stationary sounds, including speech and music along with non-stationary environmental sounds (see Table 1 for list). In the experiment, each sound was interrupted with a 2 s segment of white masking noise (Figure 2a). We asked listeners to judge whether the sound was continuous during the noise. To validate these judgments we also included trials where the sounds were unambiguously physically continuous or discontinuous during the noise (Figure 2b). We ran one version of the experiment in normal laboratory conditions, and another using online participants via Amazon’s Mechanical Turk service. The online experiment enabled the collection of data from large numbers of participants in order to better resolve differences between large numbers of stimuli.

**Figure 2.**
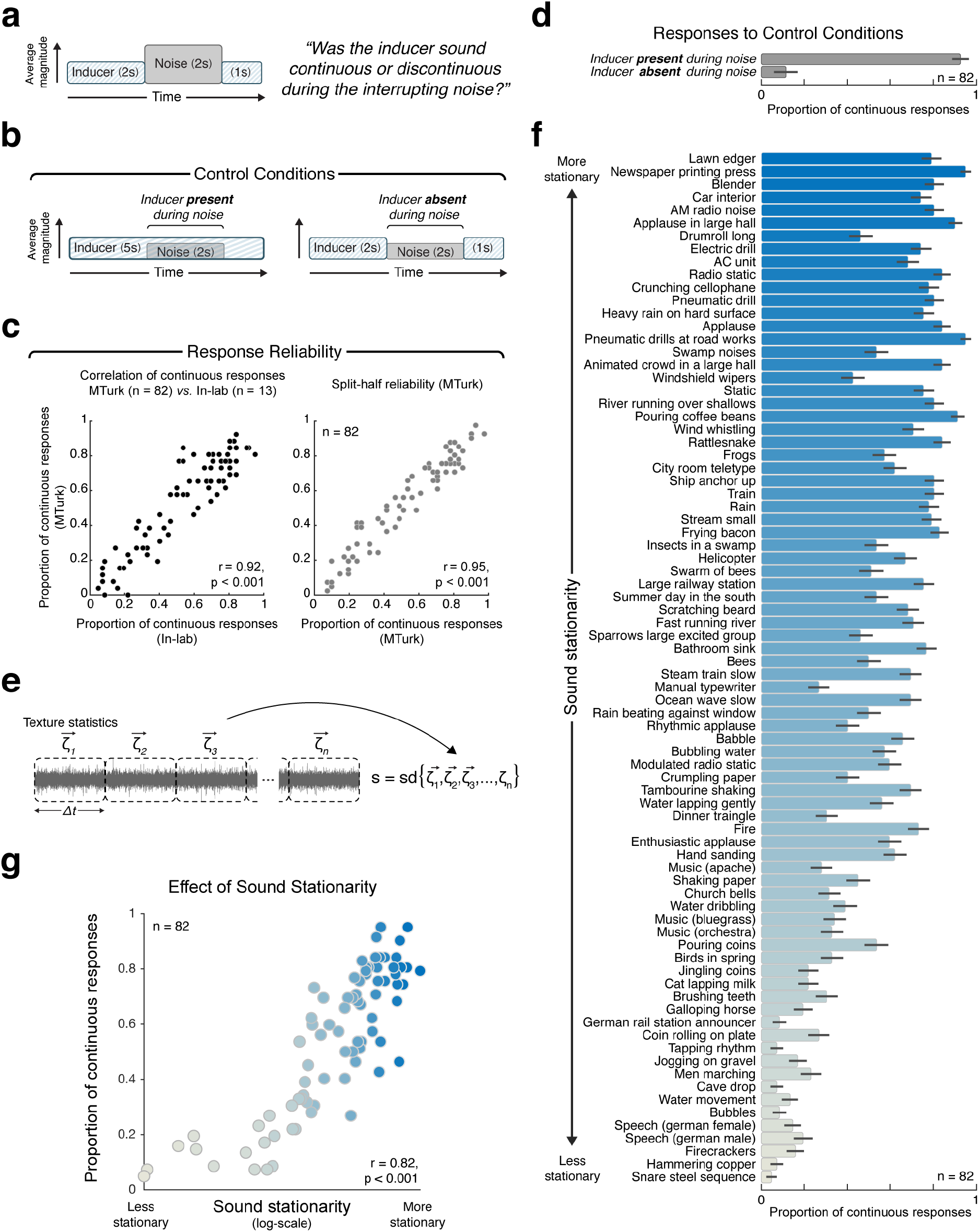
Design and Results of Experiment 1. **(a)** Schematic of stimulus and task. On each trial, one of 80 real-world sound recordings was interrupted with masking noise. The stimulus consisted of a 2 s excerpt of the real-world recording, immediately followed by 2 s of noise, immediately followed by another 1 second of the recording. Listeners reported whether the inducer sound continued during the noise. Here, and in other figures, height of inducer and noise segments in stimulus schematic symbolizes the relative sound level (in this case with the noise higher in level than the inducer). **(b)** Schematics of control condition stimuli. Left: the inducer was physically and audibly present during the interrupting noise. Right: the inducer was physically and audibly absent during the noise. In both cases the noise was 6dB lower in level (quieter) than the inducer, such that the inducer was detectably present during the noise in the left condition, and unambiguously absent during the noise in the right condition. **(c)** Reliability of results across experiments and participant splits. Left panel compares results obtained online (via Amazon’s Mechanical Turk) and in lab. Each dot plots the results for one of the 80 sounds (the proportion of participants who judged the sound to continue during the noise). The right panel shows the split-half correlation for the online experiment for one example split. The reported Pearson correlation is averaged over 10,000 random splits of participants. **(d)** Results of control conditions of Experiment 1. Here, and elsewhere, error bars show SEM. **(e)** Stationarity measure (standard deviation of statistics measured in different sound segments). Dashed lines denote borders of sound segments from which statistics were measured. The duration of the segments varied from Δ*t* = 125 ms to Δ*t* = 2 s. See Methods for details. **(f)** Results of Experiment 1 (proportion of trials on which the sound was judged to continue during the noise). Sounds are sorted by their stationarity (the value of which is denoted by the bar color). Some real world sounds have statistics that are quite stable over time (e.g. sound textures), whereas other sounds exhibit a higher degree of variability. **(g)** Mean proportion of continuous responses plotted vs. sound stationarity (also denoted by the dot color, to aid comparison to panel f).

Despite the fact that the inducer sounds were never physically present in the main condition of interest, some sounds were nearly always heard as continuing through the noise (Figure 2c). However, sounds varied in the extent to which they elicited illusory continuity: some sounds were almost never heard to continue. The results were similar between in-lab and online conditions (Figure 2c; r=0.92, p<0.001). However, the online data enabled highly reliable results across sounds, making it clear that the differences between sounds are robust and replicable (split-half reliabilities were r_MTurk_=0.95, p<0.001 and r_Inlab_=0.81, p<0.001 for the two versions of the experiment). The control conditions in which the inducer sounds were unambiguously present or absent during the noise produced high and low reports of subjective continuity, validating that listeners were correctly performing the task (Figure 2d).

To assess whether the phenomenon was specific to textures, which are distinguished by the stability of their statistics over time (i.e., stationarity), we examined whether the variation in continuity could be predicted by a sound’s temporal stationarity. We quantified stationarity as the standard deviation of a sound’s texture statistics measured in successive excerpts (Figure 2d; we computed the measure for excerpt lengths ranging from 125 ms to 2 s, and averaged the results). Sounds that were more stationary (sound textures) tended to be heard as continuous during the interrupting noise, whereas sounds that were less stationary (e.g. speech / event sounds) did not (Figure 2e). Continuity was predicted fairly well by this measure of estimated stationarity (r=0.82, p<0.001; Figure 2f).

### Experiments 2 and 3: Multi-second illusory continuity is specific to textures

The results of Experiment 1 suggest that multi-second perceptual completion could be specific to texture. However, given that the statistical structure of textures is highly predictable from one second to the next, it also seemed plausible that the extent of completion could instead be determined by the extent to which the perceptually important properties of a sound were predictable. To more thoroughly document the temporal extent of perceptual completion for different sounds, we conducted two follow-up experiments.

In Experiment 2, we varied the duration of segments of white masking noise from short (125ms) to long (2s) and measured illusory continuity for a range of “inducer” sounds (Figure 3a). Stimuli were 6 s long. The first 2 s was always a segment of an inducer sound, while the last 4 s interleaved 2 s of noise and 2 s of the inducer. For the short masker durations the noise occurred as a series of pulses interspersed with the inducer. For the longest masker duration there was a single segment of noise followed by the inducer. Listeners again reported whether they heard a sound as continuous or discontinuous during the interrupting noise. Given the large number of conditions, this experiment was conducted online (using Amazon MechanicalTurk) to obtain reliable results.

**Figure 3.**
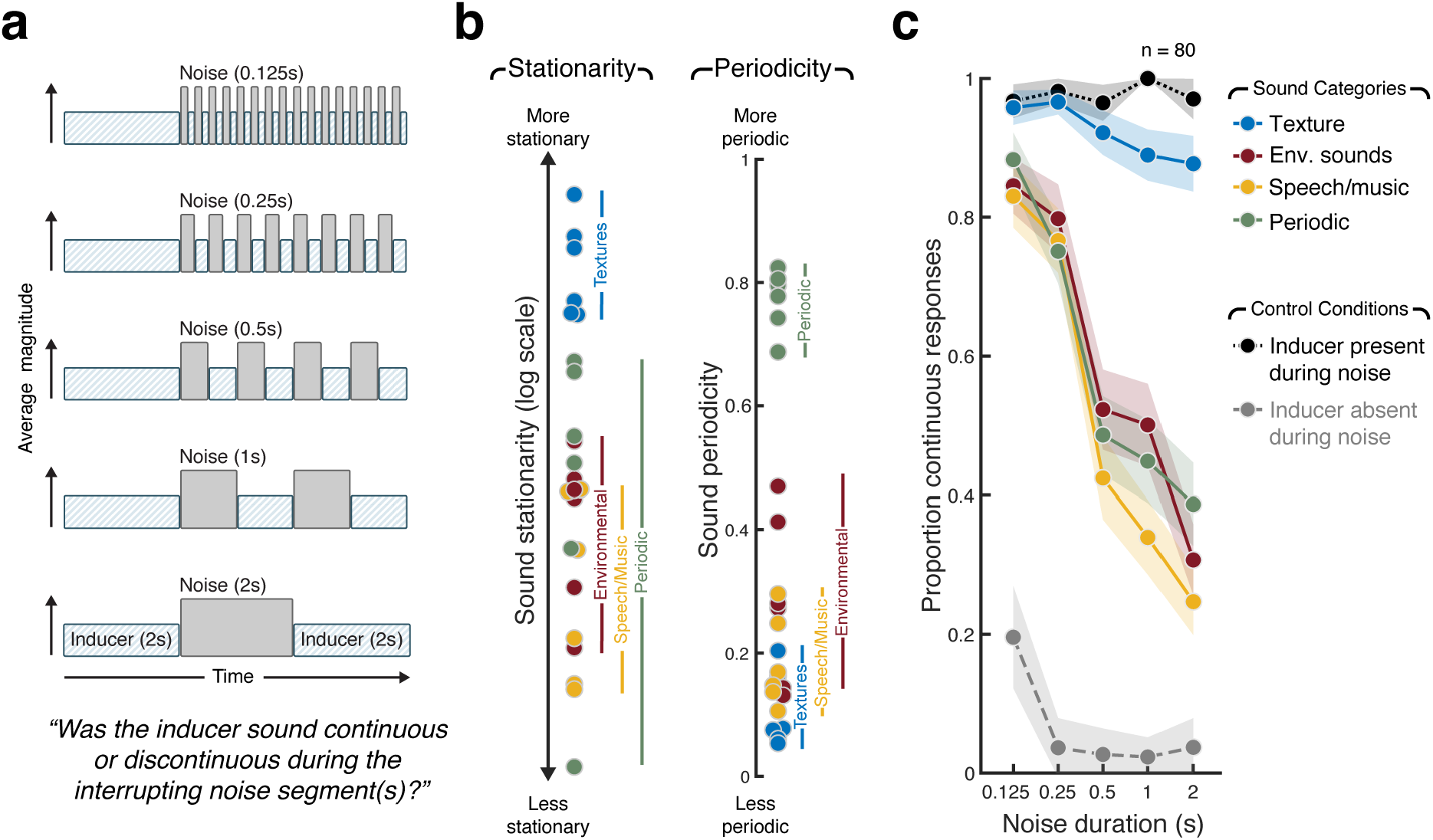
Design and Results of Experiment 2. **(a)** Schematic of stimulus conditions in Experiment 2. **(b)** Stationarity and periodicity of sounds used in the experiment. Each data point plots the stationarity or periodicity of an individual real-world sound recording used in Experiment 2 (dot color denotes the four sound categories). **(c)** Results for Experiment 2. Error bars show SEM. Control conditions show results when the inducer was physically present or absent during the interrupting noise segment(s) (same as control conditions of Experiment 1; see Figure 2b for schematics).

To better distinguish between the importance of stationarity and predictability, we included four classes of sounds: 1) stationary environmental sounds (sound textures), 2) non-stationary environmental sounds (waves, crumpling paper, church bells etc.), 3) speech and music, and 4) sounds with periodic modulations at rates ranging from 2 – 8 Hz (a galloping horse, sawing wood, ticking clock etc.). The salient properties of sounds in the latter condition were predictable over time scales of several seconds, but the sounds lacked the stable statistics of sound textures (because the modulations were slow relative to the windows over which stationarity was measured, such that different windows tended to yield different statistics). As shown in Figure 3b, these four groups of sounds were differentiated as desired via quantitative metrics of stationarity and periodicity.

Figure 3c shows the average results for the four sets of sounds. Consistent with previous work on phonemic restoration and tone continuity, all sound types exhibited considerable perceptual continuity for the shortest masker durations (no significant variation across sound class for the shortest duration, F_*3,237*_=*1.52*, p=0.*21*). But whereas sound textures exhibited a high degree of illusory continuity for the longer masker durations as well (no significant variation in perceived continuity of textures across masker durations, F_*4,316*_=*1.94*, p=0.*12*), the other three sets of sounds did not. In all three cases, the proportion of continuous responses declined sharply with increasing masker duration (Environmental sounds: F_*4,316*_=*27.41*, p<0.*001*; Speech/Music: F_*4,316*_=*32.67*, p<0.*001*; Periodic sounds: F_*4,316*_=*19.13*, p<0.*001*), producing an interaction between the effect of duration and sound class (F_12,948_=6.55, p<.001). In particular, there was a noticeable drop in perceptual continuity between 250 and 500 ms for the non-textures (Environmental Sounds: t(79)=4.82, p<0.001; Speech/Music: t(79)=4.84, p<0.001; Periodic sounds: t(79)=2.68, p=0.0091), suggestive of a limit on the extent to which non-textures can be perceptually filled in.

The results suggest that sound textures are inferred to continue over a broad range of masker durations, but that other types of sounds do so only across shorter extents. The results also indicate that the extended perceptual completion for textures is not merely the consequence of their predictability (see Supplemental Figure 1 for an analysis of the results of Experiment 1 in terms of periodicity, which provides further support for this conclusion).

In Experiment 3 we instead asked listeners to indicate the duration over which they heard continuity, using an analogue response measure. Listeners heard 6 s of audio (Figure 4a). The first 2 s was an original sound recording – either a texture, or one of a small selection of non-stationary sounds. The last 4 s consisted of white noise that could plausibly mask the original sound. Listeners reported the extent to which the original sound continued into the noise by adjusting a slider on a graphical interface once the sound had finished. The slider covered the range from 2 s (when the noise started) to 6s (when the noise ended). To confirm that participants could accurately perform this task we included control trials in which the texture was higher in level relative to the noise and physically continued either 0, 1, 2, 3, or 4 s into the noise (Figure 4b). As shown in Figure 4c, listeners positioned the slider accurately for inducer sounds that physically extended into the noise.

**Figure 4.**
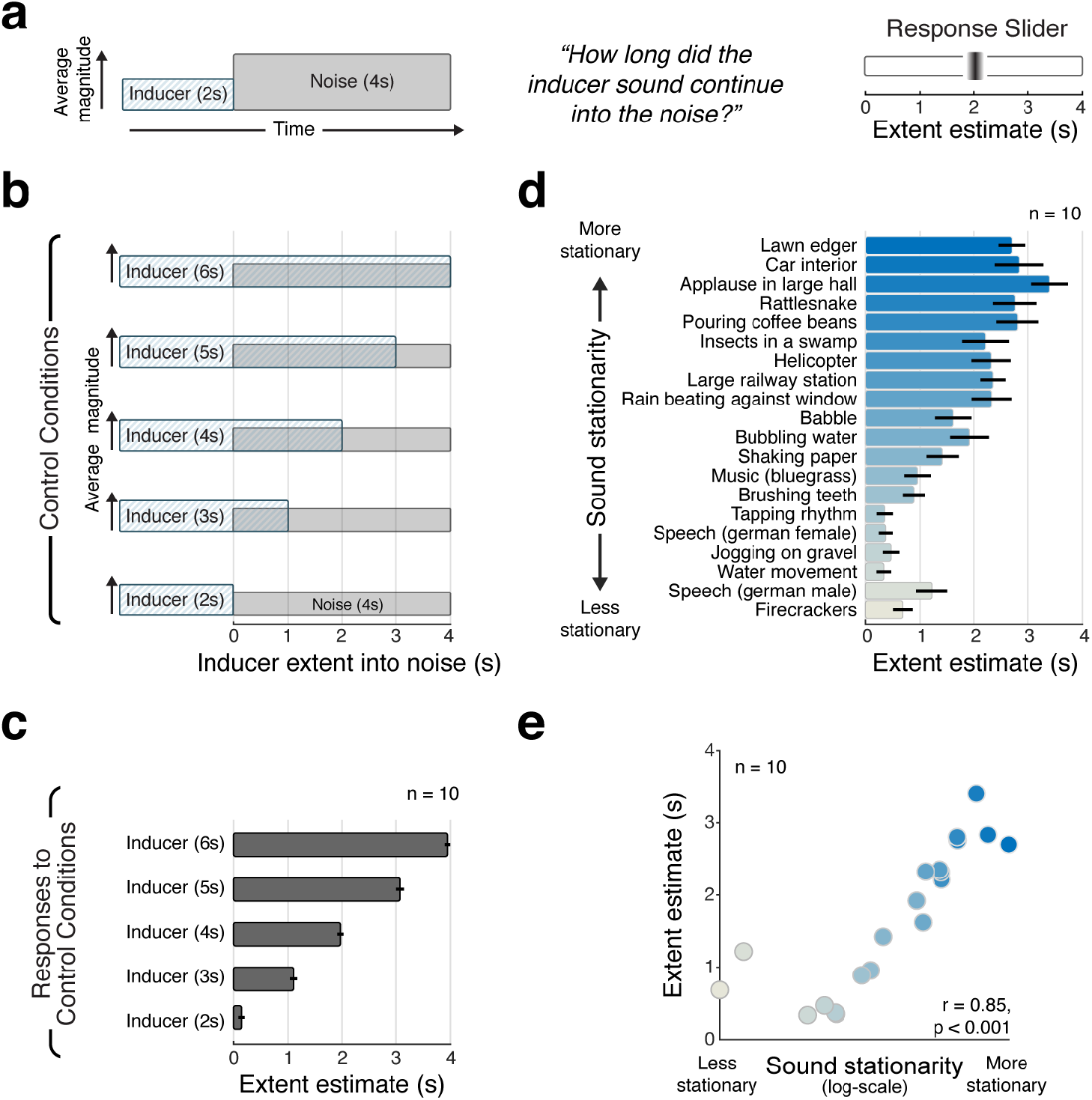
Design and Results of Experiment 3. **(a)** Schematic of stimulus and task for main condition of Experiment 3. Inducer sounds used in the experiment were real-world sound recordings. Following the noise, participants adjusted a slider to indicate the extent over which they heard the inducer sound to continue into the noise. **(b)** Schematic of stimuli for control conditions, in which the inducer was higher in level than the noise and persisted for different amounts of time after the noise onset. **(c)** Mean estimated persistence for control conditions, pooled across inducer sounds. Here and elsewhere, error bars show SEM. **(d)** Mean estimated extent of persistence for main condition of Experiment 3, plotted separately for different inducer sounds. Sounds are sorted by their stationarity. Bar color denotes inducer sound stationarity. **(e)** Mean extent estimate plotted vs. inducer sound stationarity (also denoted by the dot color, to aid comparison to panel d).

When the inducer sound was physically absent during the noise, the mean slider setting generally exceeded 2 s for texture sounds (Figure 4d, top of results graph), though it never reached all the way to 4 s. By contrast, the mean settings for non-stationary sounds (Figure 4d, bottom of graph) were largely under 1s. The sounds thus significantly varied in the temporal extent of perceived continuity (F_1,9_=90.02, p<0.001), with the extent of continuity being correlated with stationarity (Figure 4e, r=0.82, p<.001). The results suggest that there are limits to the extent of illusory continuity for textures, but provide additional evidence that they can be heard to continue substantially longer than non-textures.

### Experiment 4: Illusory texture is heard only when the texture could plausibly be masked

We next tested whether illusory texture could be explained as the inference of a masked source. We first measured whether the illusion depended on whether the interrupting noise was high enough in level to mask the texture, were they present concurrently.

Listeners performed two tasks with related stimuli (Figure 5a). In the first task, we measured the detection of a 2 s target texture superimposed on white noise, as a function of signal-to-noise ratio (SNR). On each trial listeners heard two noise bursts and were asked to identify if the target texture was present in the first or second interval. Trials were grouped into blocks related to a particular target texture. In the second task, listeners heard a 5 s texture interrupted by 2 s of white noise, and reported whether the texture was continuous or discontinuous during the noise. In both experiments, the level of the texture relative to the noise (SNR) was varied from −18 dB to +12 dB. We sought to determine whether illusory continuity would be perceived only at SNRs that would produce masking of the texture were it actually present during the noise.

**Figure 5.**
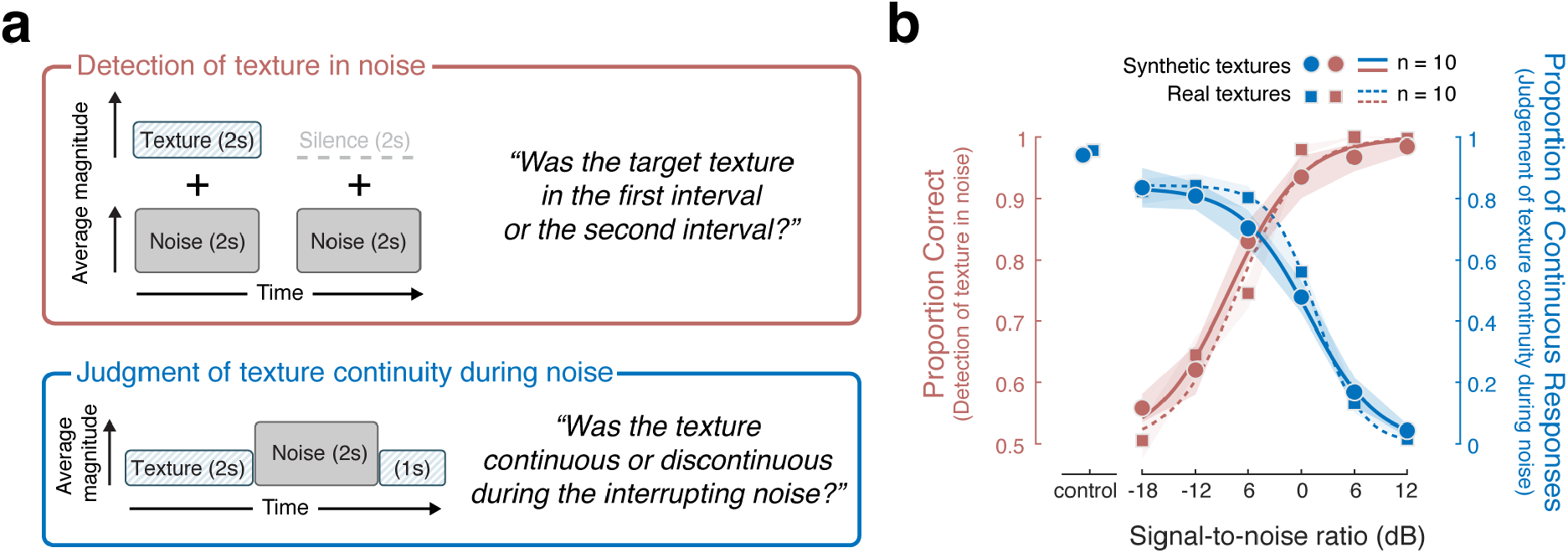
Design and Results of Experiment 4a and 4b. **(a)** Top: Detection of texture in noise (red; Experiment 4a). The stimulus consisted of two Gaussian noise signals that were 2 seconds in duration, separated by a 400 ms inter-stimulus interval. One of the signals had a texture excerpt superimposed on the noise. Listeners identified the stimulus interval that contained the texture. Bottom: Judgment of texture continuity during noise (blue; Experiment 4b). Stimulus construction and task were identical to that of Experiment 1. The stimulus consisted of a 2 s excerpt of a texture, immediately followed by 2 s of noise, immediately followed by another 1 s of the texture. We ran two versions of each experiment: one with real-world texture recordings, and one with synthetic textures generated from statistics measured from real-world recordings. Listeners judged whether the inducer texture was continuous or discontinuous during the interrupting noise segment. The SNR was varied across the same range in the two experiments. **(b**) Results of Experiments 4a and 4b. In Experiment 4a (red), the detectability of the target texture in noise decreased with SNR. In Experiment 4b (blue), listeners more readily reported the texture as continuous with decreasing SNR. The two lines correspond to the two versions of the experiments (circles and solid lines: synthetic textures; squares and dashed lines: real-world texture recordings). Shaded regions show standard error of the mean (SEM) of the individual data points. Control conditions featured stimuli where the inducer was higher in level and physically present during the interrupting noise (same as one of the control conditions in Experiment 1; see Figure 2b, left panel).

We conducted separate experiments with real-world recorded textures and synthetic textures generated from their statistics. Because the synthetic textures were generated from statistics, they additionally served to substantiate whether the perceptual completion underlying the illusory texture was statistical in nature.

As expected, detection of the target improved with increasing signal-to-noise ratio, for both real-world and synthetic textures (Figure 5b – red curves, real F_5,45_=139.9, p<0.001, synthetic F_5,45_=62.2, p<0.001). In the companion continuity task, listeners were more likely to report the inducer texture as continuous during an interrupting noise segment when the noise level was high (Figure 5b – blue curves, real F_5,45_=173.9, p<0.001, synthetic F_5,45_=91.8, p<0.001). Results were again similar for real-world and synthetic textures. Comparison of the masking curves (red) and continuity curves (blue) shows that perceived continuity was high only when listeners were poor at detecting the texture when it was physically present in the noise. The detectability and continuity of a texture were thus negatively correlated across SNRs (synthetic textures: r=−0.88, p=0.021; recorded textures: r=−0.83, p=0.040).

The SNR at which the noise masked the target varied across sounds, presumably related to some acoustic characteristic of the target sound texture (e.g. the extent of amplitude modulation; see Supplementary Figure 2 for results for individual textures). However, the SNR at which continuity was experienced also varied somewhat across sounds, and the masking thresholds and continuity thresholds were significantly correlated across sounds (r=0.7, p<0.001).

### Experiment 5: Illusory texture is eliminated by temporal gaps between texture and noise

To further test whether illusory continuity was linked to whether texture could plausibly continue during masking sounds, we measured the effect of short gaps placed between the texture and the masking noise. Listeners were presented with 5 s excerpts of textures with intervening segments of white noise (Figure 6a). The noise either abutted the texture or was temporally offset by a brief 200 ms gap on either or both sides. In all cases the noise was substantially higher in level than the texture excerpts, set individually to mask each texture based on a pilot version of Experiment 5 (see Methods). Listeners were asked to report what they heard during the noise by choosing one of six possible response contours indicating the presence of (illusory) texture over time: continuous throughout, fade-out, present only at beginning and end of the noise (‘dip’), present only in the middle of the noise (‘glimpse’), fade-in, and absent throughout (Figure 6b). We again conducted separate experiments with real-world recorded textures and synthetic textures generated from their statistics. Here we present the results with synthetic textures; see Supplementary Figure 3 for similar results with real-world recordings.

**Figure 6.**
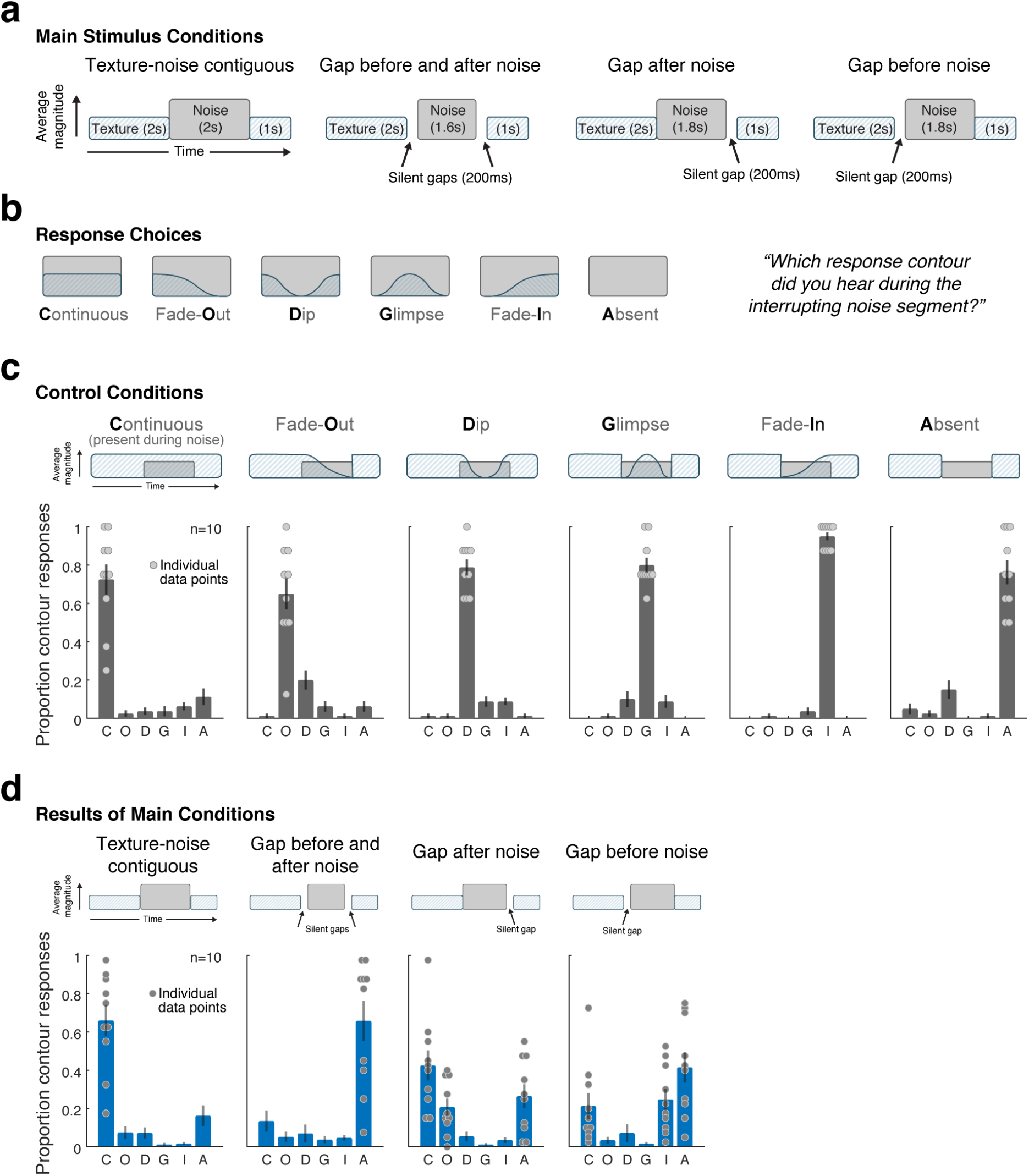
Design and Results of Experiment 5. **(a)** Listeners heard a synthetic inducer texture interrupted with masking noise and reported their perceptual experience during the interrupting noise segment. The experiment included 4 conditions, differing in the contiguity of the texture and noise (via silent gaps inserted before and/or after the noise). See Supplementary Figure 3 for analogous experiment with real-world texture recordings (which yielded similar results). **b)** Listeners chose one of six response contours to describe their perceptual experience during the interrupting noise segment. The contour response code is indicated as the bold letter for each response (e.g. “C” for “**C**ontinuous”). **(c)** To confirm task comprehension/compliance, the experiment included control trials where the texture was physically present during the intermediate noise segment and amplitude modulated according to one of the response contours. The stimulus for each condition is schematized above each of the six subplots. Graphs plot the proportion of trials on which each response was chosen. Here and elsewhere the error bars show SEM. Data for individual participants is plotted as dots for the response choices selected above chance levels for each condition. **(d)** Results of main experimental conditions. Each subplot corresponds to a condition (shown schematically above). Data for individual participants is plotted as dots for the response choices selected above chance levels for each condition.

To ensure that listeners would accurately report each of these percepts were they to occur, the experiment also included control trials in which the initial 2 s of texture was 6dB higher in level than the noise (making the texture audible when superimposed on the noise) and where the texture was modulated in amplitude to follow the contour of one of the six response choices (Figure 6b). For instance, for the “dip” control, the texture remained physically present for a short period into the noise, then faded out, then physically faded back in prior to the end of the noise. The results for these control conditions indicate that listeners accurately reported each of the six percepts when they were unambiguously present in the stimulus (Figure 6c; listeners chose the correct response contour well above chance in all cases; t(9)>8.29, p<0.001 in each condition).

As shown in Figure 6d, when the noise was contiguous with the texture, listeners predominantly reported the texture as continuing throughout the noise (t-tests comparing the proportion of continuous responses to each of the other responses were significant in each case, t(9)>4.1, p<0.01). However, silent gaps substantially altered the illusory texture. When gaps were inserted both before and after the noise, the percept of texture during the noise was largely eliminated (listeners predominantly reported the texture as being absent throughout; t-tests comparing the proportion of absent responses to each of the other responses were significant in each case, t(9)>3.51, p<0.01). A gap after the masker did not on its own eliminate illusory texture (the proportion of continuous responses was still far above chance; t(9)=6.08, p<0.001). However, the gap increased the tendency of the illusory texture to fade out before the end of the noise (t-test comparing the proportion of fade-out responses in the contiguous and gap-after-noise conditions, t(9)=3.97, p=0.0032). By contrast, a gap before the masker most often eliminated the illusory texture percept (t-test comparing the proportion of continuous responses in the contiguous and gap-before-noise conditions, t(9)=4.98, p<0.001), but illusory texture was heard to fade in more often than when gaps were both before and after the noise (t-test comparing the proportion of fade-in responses in the gap-before-and-after and gap-before-noise conditions, t(9)=3.40, p=0.0078). The latter effect suggests some degree of “retrospective” filling in^22^ driven by the texture occurring after the noise.

Together, Experiments 4 and 5 verify that textures can be heard during a few seconds of interrupting noise, even when the textures are physically absent from the stimulus. Illusory textures are heard during interrupting noise so long as the texture could have physically continued through the noise, suggesting that the illusory texture is the result of an inference about what was likely to be present during the noise. In particular, the experiments indicate that the effect is not simply due to listeners being able to imagine sounds at will during noise, as brief gaps were sufficient to largely eliminate the effect.

The results of Experiments 4 and 5 also indicate that illusory texture occurs for both for real-world recordings of texture, and for synthetic textures defined only by time-averaged statistics. The latter finding supports the idea that the illusion is the result of extrapolated statistical properties.

### Experiment 6: Illusory sound texture is represented similarly to actual sound texture

What happens in the auditory system to produce illusory texture? To probe the relationship between the representation of illusory texture and texture that is physically present in the sound signal, we sought to leverage prior findings that listeners’ texture judgments are biased by several seconds of stimulus history^20^, presumably reflecting the averaging process underlying texture statistics. If illusory texture is represented like actual texture, for instance with persistent activity in the relevant part of the auditory system, it might similarly bias judgments of subsequent texture.

We modified a texture discrimination experiment from our previous work^20^ to incorporate illusory texture. The experiment required listeners to judge which of two sound texture excerpts was most similar to a reference texture. The two excerpts were generated from statistics lying on a line between the mean statistics of a large set of textures and the reference texture. The first excerpt was 5 s and had a subtle change (“step”) in statistics at some point during its duration (Figure 7a). The second excerpt (the “morph”) was 2 s and had constant statistics at discrete points sampled on a line between the mean and the reference texture. Listeners were informed that the first excerpt would undergo a change at some point, and they should base their judgments on the sound of the texture at its endpoint. We previously found that these texture steps biased listeners’ judgments away from the endpoint if they occurred within a few seconds of the endpoint, suggestive of an integration window of several seconds for estimating sound texture statistics. We tested whether illusory texture heard during an interrupting noise burst would similarly bias texture judgments.

**Figure 7.**
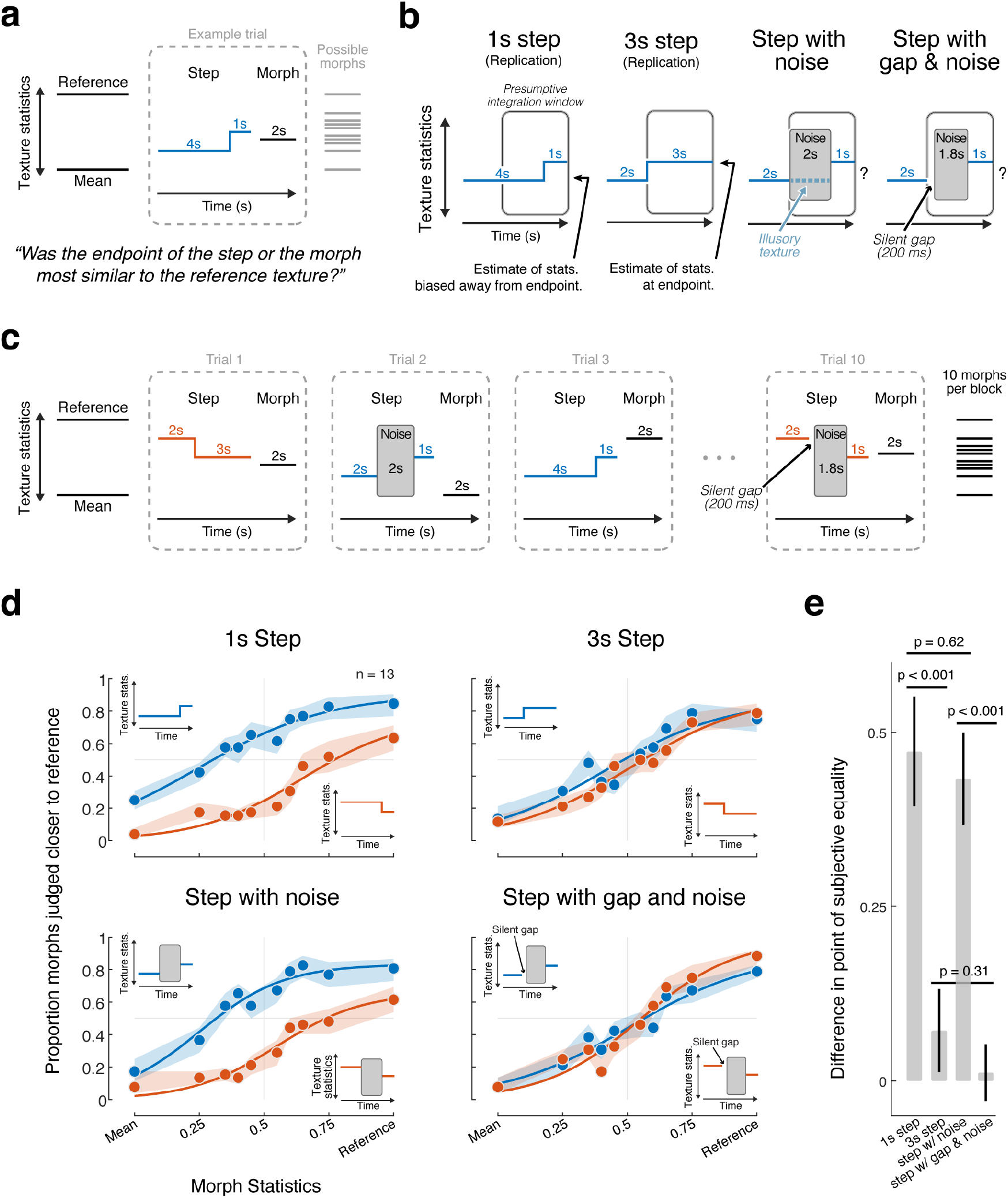
Design and Results of Experiment 6. **(a)** Step experiment paradigm. At the start of each trial block, participants were familiarized with a reference texture (synthesized from the statistics of a real-world texture recording) and a mean texture (synthesized from the average statistics of 50 real-world texture recordings). On each trial participants heard two stimuli and judged which stimulus was most similar to the reference, basing their judgments on the endpoint of each stimulus. The first stimulus contained a subtle “step” in statistics, the direction and temporal position of which varied from trial to trial. The morph had constant statistics in a given trial, but varied between the reference and mean across trials. **(b)** Main conditions and interpretation. Previous experiments suggested that humans estimate texture statistics using a temporal integration window of several seconds (symbolized in grey). A step occurring 1 second from the endpoint (first panel) is thus predicted to bias judgments away from the endpoint. By contrast, a step occurring 3 seconds form the endpoint (second panel) should yield little or no bias in judgment away from the endpoint. In the critical condition (third panel), the middle of the step stimulus was replaced with 2 s of Gaussian masking noise. This was expected to elicit an illusory texture during the presumptive integration window that could bias judgments away from the endpoint if the illusory texture replicated the effects of an actual texture in the relevant part of the auditory system. The last condition contained a 200 ms silent gap before the noise. This was a control condition to confirm that eliminating the percept of illusory texture would eliminate any bias observed in the third condition. **(c)** Trial blocking. The experiment was divided into blocks of 10 trials related to a particular reference texture. The step condition and direction (towards or away from the reference) varied from trial to trial (randomly ordered over the course of the experiment subject to the blocking constraint). The 10 morph positions were presented once per block in random order. **(d)** Results of Experiment 6. Here and elsewhere, shaded regions show SEM of individual data points obtained by bootstrap and curves plot logistic function fits. **(e)** Plot of the bias produced by the step in each condition, quantified as the difference between points of subjective equality for the upward and downward step conditions. Error bars show SEM, obtained by bootstrap (10,000 samples).

In the experiment, the first interval always contained a change in statistics, either at 1 s from the endpoint or 3 s from the endpoint (Figure 7b). Based on our prior work, we expected the 1 s step to bias discrimination judgments, but for the bias to be greatly reduced for the 3 s step, as it occurs outside the apparent integration window for texture^20^. The two critical conditions included interrupting white noise (Figure 7b). In one condition, the noise extended from 3 s to 1 s from the endpoint. Based on Experiment 5b, we expected illusory texture to be heard during the noise, extrapolated from the first part of the step. In another condition the first 200 ms of the noise was replaced by a silent gap, which we expected to eliminate the illusory texture. The gap also served as a control for the possibility that the noise on its own might bias texture perception. Trials from the different conditions were intermixed within blocks corresponding to a particular reference texture (Figure 7c), and in all cases listeners were instructed to base their judgments on the end of the step stimulus. If the perception of illusory texture instantiates representations in the auditory system like those of physically realized texture, the step with noise, but not the step with gap and noise, might produce a bias in listeners’ judgments.

As shown in Figure 7d, listeners were biased by texture steps that occurred at 1 s before the endpoint but not at 3 s before the endpoint, as expected. This effect was quantified by fitting psychometric functions to the data and measuring the bias as the difference in the point of subjective equality between the two functions (Figure 7e). The bias was significantly different for the 1 s step and 3 s step conditions (p<0.001, via bootstrap), and not significantly different from 0 for the 3 s step (p=0.22). But when the step was interrupted by noise, listeners exhibited a bias comparable to the 1 s step condition, as though the statistics of the initial texture segment were heard during the noise and incorporated into the estimate for the subsequent texture segment (no significant difference in bias between 1 s step and noise conditions, p=0.62). The bias was eliminated by the silent gap preceding the noise (significantly larger bias for noise than gap condition, p<0.001; no difference between 3s step and gap conditions, p=0.31; and the bias for the gap condition was not significantly different from 0; p=0.77).

The results indicate that illusory texture biases texture judgments similarly to texture that is physically present. The effect provides objective evidence for the presence of illusory texture, and places constraints on its representation in the auditory system. In particular, the results suggest that the illusion involves changes to the representations that are integrated to form texture statistic estimates, as these estimates appear to be altered by the illusion.

### Experiment 7: Textures are inferred to continue even when concurrent with non-textures in auditory scenes

In the final experiment we probed the real-world relevance of illusory texture by exploring filling in for typical auditory scenes, which might consist of multiple sound sources that vary in stationarity. We designed an experiment in which listeners were again asked to report whether a sound was continuous or discontinuous during a segment of interfering noise. However, in some conditions two inducers were presented simultaneously to form a simple auditory scene (Figure 8a). The inducer pairs consisted of one texture and one non-texture sound, selected based on their stationarity (Figure 8b). On each trial listeners heard the auditory scene (sound pair) interrupted by noise: 2 s of the inducer pair followed by 2 s of interrupting white noise and then another 1 s of the inducer pair. Prior to the auditory scene the listeners were cued with a 2 s excerpt of the target sound whose continuity they were supposed to report. The cued sound could be the texture, the non-texture, or the sound pair.

**Figure 8.**
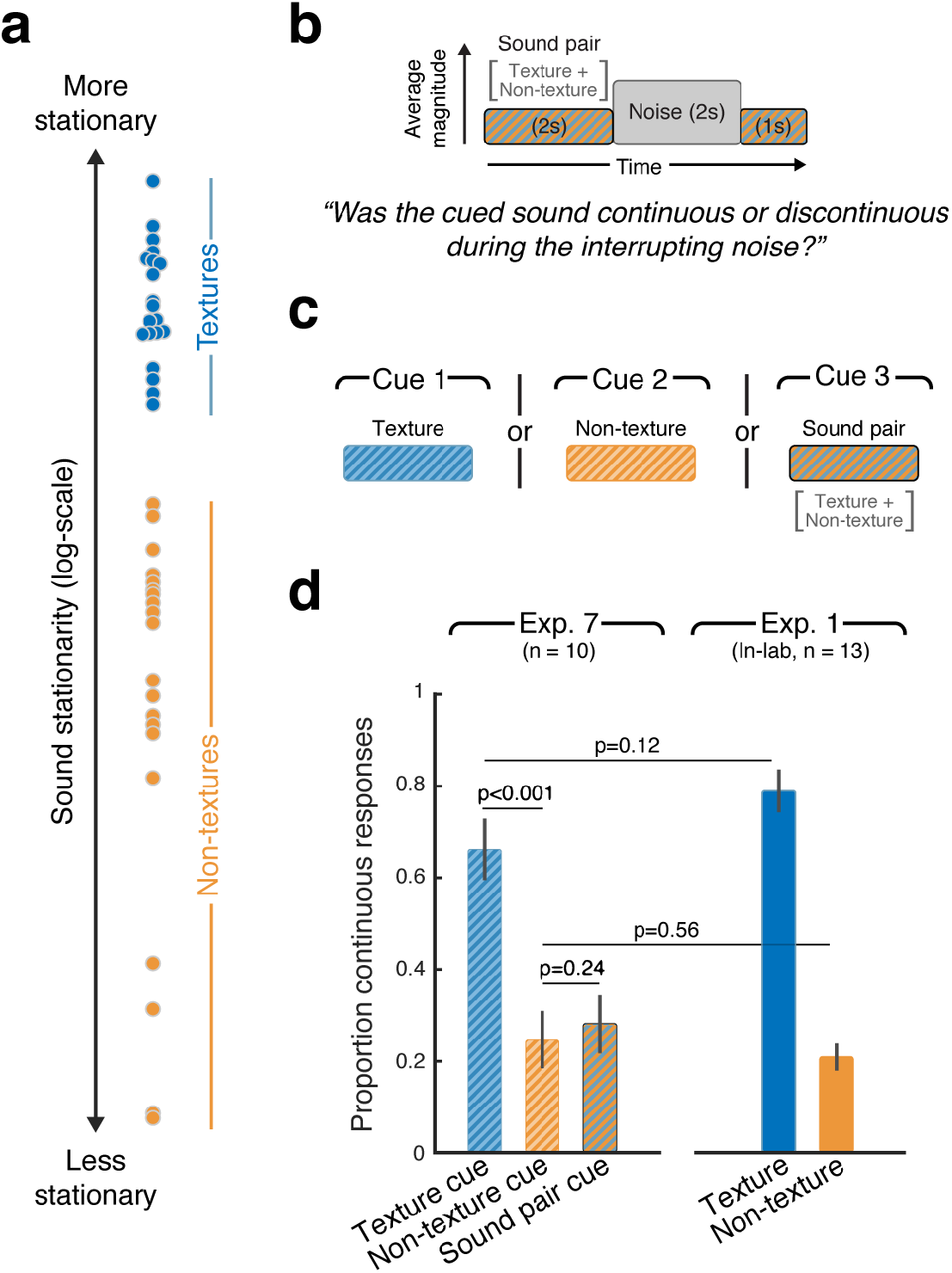
Design and Results of Experiment 7. **(a)** Stationarity of the 40 real-world sound recordings used as inducers in Experiment 7. The sounds were divided into two groups: 20 textures and 20 non-textures (selected based on their sound stationarity). **(b)** Schematic of the main stimulus conditions in Experiment 7. The inducer sound was the superposition of a texture with a non-texture (sound pair), and was interrupted by masking noise as in previous experiments. **(c)** A cue sound was presented to the listeners prior to each trial, indicating which sound they should be judging the continuity of. The cue could either be the texture, the non-texture or the superposition of the two. **(d)** Results of Experiment 7, plotting the proportion of continuous responses for each of the three conditions. Error bars show SEM. Right bars: The results of the same task for the 20 textures and 20 non-textures presented in isolation, replotted from Experiment 1.

Even though both sounds were present prior to the noise, only the texture sound was heard during the noise to a substantial extent (Figure 8c; continuity was greater for the texture than for the non-texture, t(9)=9.97, p<0.001, or the scene, t(9)=8.09, p<0.001. The proportion of trials on which the texture and non-texture were reported to continue was similar to when the sounds were present in isolation (Figure 8c, replotting data from Experiment 1; no significant difference in continuity for textures in isolation vs. paired with a non-texture: t(21)=1.61, p=0.12; or for non-textures: t(21)=0.59, p=0.56). This result suggests that perceptual completion is determined at the level of individual streams, and that textures are streamed separately from other sounds. It also suggests that the presence of non-stationary sounds does not prevent extended completion of concurrent textures.

## Discussion

We documented an illusion in which sound textures are perceived to continue over periods of up to several seconds when interrupted by a foreground sound, despite being physically absent. This “illusory texture” had a number of diagnostic characteristics. First, the temporal persistence of the illusion appears to be related to the temporal stationarity of the sound. Illusory sound texture lasted up to several seconds during extended masking noise, whereas temporally variable sounds, such as speech, faded out after a few hundred milliseconds. This difference appears to not be simply explained by the timescale over which signals are predictable, because temporally sparse periodic sounds (regular rhythms etc., that are highly predictable) also failed to produce multi-second illusory continuity. Extended illusory continuity may thus be unique to statistical representations of sound. Second, illusory texture only occurred when the interrupting foreground sound was temporally contiguous with the texture and sufficiently intense as to plausibly mask the texture. These two results suggest that illusory texture is the result of an inference about whether the texture continues during the masking noise, analogous to that classically thought to occur for tones or speech. Third, leveraging the multi-second averaging process that underlies texture perception, we found that illusory sound texture was incorporated into subsequent judgments similarly to actual sound texture. This finding provides objective evidence for the illusion and suggests that it could be mediated by persistent activity within the same representational locus as actual texture. Fourth, when a texture co-occurred with temporally variable sounds, as in typical auditory scenes, the texture was perceived to continue during an interrupting masker even though comparable illusory continuity did not occur for the nonstationary concurrent sounds. This observation suggests that textures are represented as streams within auditory scenes, and that perceptual inferences of continuity are made on individual streams. Given the ubiquity of texture, and of concurrent foreground sounds, the results suggest that the background of auditory scenes is filled in routinely without the listener realizing it, constructing a stable representation of the auditory world from impoverished sensory data.

### Comparison to previously described continuity illusions

Perceptual filling in of speech and tones, and its relation to masking, was first described many decades ago^3–6^. Illusory texture differs from these previously described perceptual completion effects in at least two respects. First, the duration of the effect is much longer. Tones and speech are heard to complete only over a few hundred milliseconds^23–25^. We documented this difference in Experiments 1-3, finding that textures are unusual among natural sounds in persisting for long durations. To our knowledge the only precedent for this observation is an informal note by Warren^6^ that synthetic noise could be heard to continue for relatively long periods of time when interrupted by another masking noise, though this was apparently never substantiated experimentally. The extent of continuity may relate to the likelihood that textures are continuous in the world. Textures are often generated by large numbers of concurrent events (raindrops, hand claps, insect noises), potentially making them unlikely to stop abruptly (unlike, for instance, a person talking). The extent of illusory continuity for different sounds could thus be a rational inference about the likely state of sounds in the world.

It is natural to wonder whether the extent of illusory continuity might alternatively relate to integration windows for texture statistics, which appear to be several seconds in duration^20^, on par with the extent of illusory texture. One reason to think that this similarity of time scales is coincidental is that less stationary textures appear to be integrated over longer periods of time, perhaps as needed to obtain stable statistic estimates^20^. By contrast, the extent of illusory continuity is, if anything, longer for more stationary textures (Experiment 3; Figure 4d). It thus seems more plausible that the extent of perceptual completion reflects priors on the continuity of different types of sound sources.

A second difference with classical continuity effects is that the filling in of texture is statistical. Unlike tones or speech, textures are stochastic, and on the time scales at which the continuity occurs can only be described and extrapolated in terms of the distribution of their features. There are some previously documented examples of perceptual completion of abstracted properties of sound, in that frequency and amplitude modulation appear to be filled in at a level of representation that discards phase^26,27^. Texture completion appears to be a more extreme version of this sort of phenomenon, in that the content that is completed reflects time-averaged properties of the inducing stimulus.

The statistical nature of illusory texture provides additional evidence for statistical representations of texture. Previous evidence for statistical representations came from the recognizability of textures synthesized from statistics^14,15,18^, or from the discrimination of texture excerpts^16^. But for texture that is physically realized in a sound signal, any statistical representation is concurrent with, and presumably derived from, the representation of the acoustic details composing a texture. As a result, the role of statistical representations in perception must be inferred indirectly. Illusory texture, by contrast, occurs without the presence of any underlying acoustic elements, and when the inducing texture is defined only by statistics, the illusory texture that is heard during the noise must be mediated exclusively by a completed statistical representation. In that sense it provides the clearest evidence yet that texture perception is based on statistics. We also note that the extrapolated statistics could nonetheless instantiate a representation of illusory acoustic details that are consistent with the extrapolated statistics (Experiment 6 is consistent with this possibility).

### Neural basis of illusory texture

Experiments with traditional continuity illusions have in some cases suggested explicitly “filled in” neural representations, envisioned as persistent activity from the inducer stimulus continuing during the noise. Evidence for persistent activity has been seen in primary auditory cortex of cats and monkeys for the original tone continuity illusion^28,29^. And several studies in humans have reported evidence of explicit filling in during phonemic restoration^30–32^. On the other hand, other studies of tone continuity have suggested more implicit representations of the continuity, with the tone onsets and offsets being suppressed by the noise^33–35^, and one report that illusory tones do not produce aftereffects comparable to those induced by real tones^36^.

Our results are suggestive of an explicit representation of illusory texture. We found evidence that illusory texture biases the perception of subsequent texture, as though the illusory texture was incorporated into the integration window for estimating texture statistics (Experiment 6). This result is consistent with the idea that perceptual continuity is mediated by persistent activity at the representational stage that is averaged to yield texture statistics. Illusory texture could thus involve a conceptually similar mechanism to that proposed for tones, though in a higher-order representation (e.g. modulation filters^37,38^), and with persistent activity lasting much longer than with tones. But given that the continuity appears to be driven by the statistical properties of a sound, and that these statistics are derived via a multi-second integration process^20^, any persistent activity may be driven by feedback from a representation of those statistics, the neural locus of which remains unclear. The strength and duration of the illusion should enable neurophysiological experiments to more definitively probe its neural basis.

### The role of subjective perception

As with other auditory continuity effects, illusory texture is heard in conditions where the texture would be masked were it physically present. Sounds are thus subjectively audible in conditions where they would be undetectable. This subjective audibility is particularly apparent for illusory texture because of how long it lasts, creating vivid percepts that facilitate introspection. The subjective audibility of masked sounds appears to differentiate auditory continuity from perceptual completion effects in vision. The conditions producing auditory continuity (masking) are most analogous to those eliciting “amodal” completion in vision, in which occluded contours (that are obscured by foreground objects) are inferred to extend behind the foreground object (Figure 1a). Amodal completion has measurable consequences for perception, for instance constraining face recognition^39^ and motion integration^40^, but the completed contours are not “seen” in the way that perceptually completed sounds are “heard”. The subjective percept of auditory continuity is in this respect more similar to “modal” completion, in which part of a foreground object that has low contrast with the background is nonetheless subjectively visible (Figure 1a). This difference may reflect the fact that there are two different physical situations to distinguish in vision (occlusion, in which the occluded object cannot possibly be seen, and camouflage, in which the foreground object is invisible only because of accidental matches with the background color). Seen contours appear to be a code for occluding objects. This distinction is absent in audition, because sounds superimpose rather than occlude. Subjective audibility thus appears to be used to code for the presence of sound sources even in conditions where they are physically undetectable and must be inferred.

## Methods

### Auditory Texture Model

The auditory texture model processed an input sound waveform via a cascade of filter banks and measured statistics of their outputs. The filter bank cascade was identical to that described in an earlier paper^20^, which was a modification of the original McDermott and Simoncelli model^14^. The first filter bank approximated the frequency selectivity of the cochlea. The envelopes of the output of these filters were then filtered with a temporal modulation filter bank. The “texture” statistics measured from these representations consisted of the mean, coefficient of variation, and skewness of the cochlear envelopes, pair-wise correlations across cochlear envelopes, the power from the modulation filters, and pair-wise correlations across modulation bands. These statistics were identical to those used previously [cite] except that the cochlear envelope kurtosis was omitted, as we have found it to have little impact on synthesis quality.

For completeness we describe the model in full here. The rest of this section is reproduced from the methods section of the original paper^20^.

To simulate cochlear frequency analysis, sounds were filtered into subbands by convolving the input with a bank of bandpass filters with different center frequencies and bandwidths. We used 4^*th*^-order gammatone filters as they closely approximate the tuning properties of human auditory filters and, as a filterbank, can be designed to be paraunitary (allowing perfect signal reconstruction via a paraconjugate filterbank). The filterbank consisted of 34 bandpass filters with center frequencies defined by the equivalent rectangular bandwidth (ERB_N_) scale^41^ (spanning 50Hz to 8097Hz). The output of the filterbank represents the first processing stage from our model (Supplemental Figure 4).

The resulting “cochlear” subbands were subsequently processed with a power-law compression (0.3) which models the non-linear behavior of the cochlea ^42^. Subband envelopes were then computed from the analytic signal (Hilbert transform), intended to approximate the transduction from the mechanical vibrations of the basilar membrane to the auditory nerve response. Lastly, the subband envelopes were downsampled to 400 Hz prior to the second processing stage.

The final processing stage filtered each cochlear envelope into amplitude modulation rate subbands by convolving each envelope with a second bank of bandpass filters. The *modulation filterbank* consisted of 18 half-octave spaced bandpass filters (0.5 to 200 Hz) with constant *Q* = 2. The modulation filterbank models the selectivity of the human auditory system and is hypothesized to be a result of thalamic processing^37,43,44^. The modulation bands represent the output of the final processing stage of our auditory texture model.

The model input was a discrete time-domain waveform, *x*(*t*), usually several seconds in duration (~5 s). The texture statistics were computed on the cochlear envelope subbands, *x*_*k*_(*t*), and the modulation subbands, *b*_*k,n*_(*t*), where *k* indexes the cochlear channel and *n* indexes the modulation channel. The windowing function, *w*(*t*), obeyed the constraint that Σ_*t*_ *w*(*t*) = 1.

The envelope statistics include the mean, coefficient of variance, skewness and kurtosis, and represent the first four marginal moments. The marginal moments capture the sparsity of the time-averaged subband envelopes. The moments were defined as (in ascending order)

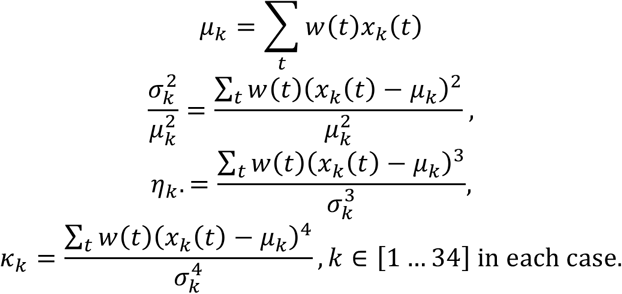

Pair-wise correlations were computed between the eight nearest cochlear bands. The correlation captures broadband events that would activate cochlear bands simultaneously^14,15^. The measure can be computed as a square of sums or in the more condensed form can be written as

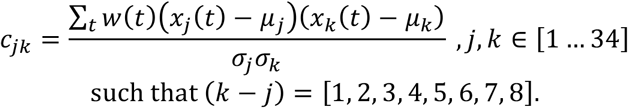

To capture the envelope power at different modulation rates, the modulation subband variance normalized by the corresponding total cochlear envelope variance was measured. The modulation power measure takes the following form

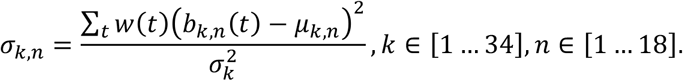

Lastly, the texture representation included correlations between modulation subbands of distinct cochlear channels. Some sounds feature correlations across many modulation subbands (e.g. fire), whereas others have correlations only between a subset of modulation subbands (ocean waves and wind, for instance, exhibit correlated modulation at slow but not high rates^14^). These correlations are given by

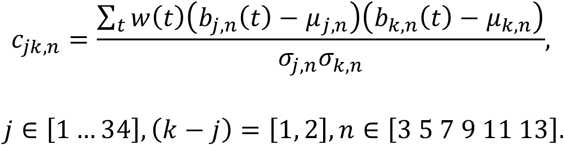

The texture statistics identified here resulted in a parameter vector, ζ, which was used to generated the synthetic textures.

### Auditory Texture Synthesis

Synthetic textures were generated using a previously published variant of the McDermott and Simoncelli^14^ synthesis system^20^. The sound texture synthesis system measured the time-averaged statistics from a real-world texture recording using the auditory texture model. The statistics were measured from 7 s sound recording excerpts. The measured statistics were subsequently imposed on the cochlear envelopes of a 7 s Gaussian noise seed via an iterative procedure that adjusted the statistics of the synthetic sound to match the statistics of the target real-world sound using the limited-memory Broyden–Fletcher–Goldfarb–Shanno (L-BFGS) gradient descent optimization algorithm^14,20^. By seeding the synthesis system with different samples of noise, the procedure allowed for the generation of distinct exemplars that possessed similar texture statistics. Code implementing the synthesis procedure is available online: https://github.com/rmcwalter/STSstep.

### Sound Stationarity Metric

To quantify the stationarity of real-world sound recordings, we computed the standard deviation of the texture statistics measured across successive segments of a signal, for a variety of segment lengths (0.125, 0.25, 0.5, 1, and 2 s). For each segment length we carried out the following steps. First, we divided a signal into adjacent segments of the specified duration and measured the statistics in each segment. Second, we computed the standard deviation across the segments. Third, we averaged the standard deviation across statistics within each statistic class (envelope mean, envelope coefficient of variation, envelope skewness, envelope correlations, modulation power, and modulation correlations). Fourth, we normalized the average standard deviation for a class by its median value across sounds (to put the statistic classes on comparable scales, and to compensate for any differences between statistics in intrinsic variability). Fifth, we averaged the normalized values across statistics classes. We then averaged the resulting measure for the five segment lengths to yield a single measure of stationarity for each real-world sound recording.

### Experimental Procedure for In-Lab Experiments

The Psychophysics Toolbox (Psychtoolbox) for MATLAB was used to play sound stimuli and collect responses. All stimuli were presented at a sampling rate of 48kHz in a soundproof booth (IAC Acoustics). The presentation sound pressure level (SPL) varied across sounds and experiments, and is specified in the sections below. Sounds were played from the sound card of a MacMini computer over Sennheiser HD280 PRO headphones.

### Experimental Procedure for Online Experiments

Some experiments necessitated small numbers of trials per participant, and thus required a large number of participants to obtain reliable results. We conducted these types of experiments using Amazon Mechanical Turk crowdsourcing platform. In previous work we have found that online data can be of comparable quality to in-lab data provided some modest steps are taken to standardize sound presentation^45^. Each participant first adjusted their volume setting so that a calibration Gaussian noise sound was at a comfortable level. The noise root-mean-squared (rms) level was set to the maximum of the rms level of the experimental stimuli. Participants then completed a ‘headphone check’ task to help ensure that they were wearing headphones or earphones^46^. Finally, to eliminate participants who were not fully focused on the task, we included practice trails at the beginning of the experiment (described in more detail below). On these trials the inducer stimulus was physically and audibly present or absent during the noise. We excluded participants who performed at less than 75% correct on these practice trials.

### Masking Noise Level

All experiments apart from Experiment 4 used a common stimulus structure in which an inducer sound was interrupted by a white Gaussian noise masker that was higher in level than the inducer. To ensure the interrupting noise would plausibly mask the inducer were they presented simultaneously, the first author (RM) performed a brief experiment to set the level of the masking noise for each inducer sound. In the experiment, 2 s of Gaussian noise was superimposed on 2 s of inducer sound. The 2 s of sound was pulsed on and off, with an inter-stimulus-interval of 400 ms. For each “on” pulse, a new 2 s noise waveform was generated and the 2 s inducer was randomly sampled from a longer 7 s excerpt. The inducer level was fixed at 55dB SPL. The level of the noise was increased or decreased, in half dB increments starting from 55dB SPL, until the noise was sufficiently high in level to mask the inducer sound. The resulting signal-to-noise ratio (SNR) between the inducer and the noise is provided for each experiment in Supplemental Tables 1–6.

### Experiment 1 – Illusory continuity for a large set of natural sounds

#### Stimuli

Eighty natural sounds were selected that spanned a wide range of source types (e.g. speech/music, textures, event-like sounds; see Supplemental Table 1 for a list of the 80 sounds used in Experiment 1) and sound stationarity (Supplemental Figure 5 shows the measured stationarity of sounds). The stimuli were generated from 5 s excerpts sampled from longer 7 s natural sound recordings (with the excerpt onset randomly sampled between 0 and 2 s). The 5 s excerpt was interrupted with white Gaussian noise, such that the stimulus consisted of the concatenation of 2 s of the real-world sound recording (the “inducer”), 2 s of Gaussian noise (the “masker”), and the remaining 1 s of the inducer. The inducer and noise were cross-faded over 20 ms using raised cosine ramps applied to the onset and offset of the two signals, to avoid abrupt phase discontinuities in the inducer. As a result, the signal excerpts used to construct the stimulus were 2.01 s, 2.02 s, and 1.01 s, with the midpoint of the crossfade occurring 2 s and 4 s into the resulting stimulus. The intermediate noise segment otherwise replaced the synthetic texture, so that the synthetic texture and the noise were not physically present at the same time. The level of the interrupting noise was set to mask the inducer were they presented simultaneously. This level was set individually for each inducer sound as described in the previous section (Supplemental Table 1).

Control trials and practice trials used stimuli that were identical to those in the experimental trials, except that a) the inducer sound level was set to +6dB relative to the noise, and b) the inducer was physically present during the intermediate noise segment on half of the trials. These trials were intended to confirm task comprehension (i.e., we expected that participants who were performing the task as desired would judge the trials with the physically present inducer as continuous, and the trials with the physically absent inducer as discontinuous). The control trials and practice trials used a distinct set of forty real-world sound recordings.

#### Procedure (In-lab)

Participants judged whether the inducer sound was continuous or discontinuous during the intermediate noise segment. Each participant performed 200 trials: two presentations for each the 80 sounds and two presentations of the 20 control trials with randomly selected inducer sounds. The presentation level of the inducer sound was set to 55dB SPL. The presentation order was randomized for each participant. The participants did not receive feedback during the main experiment.

Prior to the main experiment, participants performed 20 practice trails with feedback. The sounds used for the 20 practice trials did not overlap with the sounds used for the 20 control trials.

#### Procedure (Online)

Prior to the experiment, participants were instructed to set the volume of their headphones to a comfortable level while listening to a calibration Gaussian noise signal set to the maximum noise level that would be presented during the main experiment.

Participants then performed a brief headphone check experiment [Woods et al. 2017] intended to weed out participants who were ignoring the instructions to wear headphones or earphones. They next completed 20 practice trails in which they were asked to report whether they “heard the sound” or “didn’t hear the sound” during the interrupting noise segment. If the participant’s performance met or exceeded 75% on the practice section, they continued to the main experiment, otherwise the session was ended. At this point the calibration noise was presented a second time before the participants began the main experiment to ensure comfortable listening conditions. The main experiment consisted of 100 trials: 80 trials of the main experimental condition (one for each of the 80 sounds) and 20 control trials. The sounds used for the 20 practice trials did not overlap with the sounds used for the 20 control trials.

#### Participants

Online: 140 participants began the online experiment, of which 58 failed either the headphone check or the practice session. The remaining 82 participants completed the main MTurk experiment (40 female, mean age = 38.8, s.d. = 10.9). MTurk participants were unique to Experiment 1. In-lab: The 13 in-lab participants all exceeded 75% on the practice session and were included in the analysis (6 female, mean age = 23.5, s.d. = 2.3).

### Experiment 2 – Illusory continuity across different masker durations

#### Stimuli

The stimuli consisted of excerpts of real-world sound recordings interrupted with Gaussian noise segments. Twenty-four source recordings were used in the experiment, selected from 4 categories: speech/music, textures, periodic/rhythmic sounds, and environmental sounds (Table 2). The textures were selected from a larger set as the most stationary (according to the measure described earlier). To select the periodic sounds, we measured the normalized auto-correlation of the envelope of the broadband waveform (the Hilbert envelope of the waveform, down-sampled to 400 Hz). The height of the largest peak between 125 ms and 500 ms (2 – 8 Hz) was selected as the measure of periodicity. The six periodic/rhythmic sounds were selected from a larger set as those that had the highest values of this measure. The environmental sounds were selected to be less stationary than the textures and less periodic than the periodic/rhythmic sounds. The speech and music sounds comprised recordings of English, German, bluegrass, and classical.

To generate the stimuli we first selected a 6 s excerpt of one of the sources, where the onset was randomly selected from the first 1 s of a 7 s recording. Segments of the 6 s sample were then replaced with Gaussian white noise. The first 2 s of the sample was left intact. 50% of the remaining 4 s was replaced with noise, such that the stimulus alternated between noise and the inducer. The duration of the noise segments varied logarithmically between 125 ms and 2 s, resulting in between 16 and 1 segments (Figure 3a). The level of the noise segment(s) was set individually for each inducer to a level that would mask the inducer were the noise and inducer present concurrently (see Table 2 for levels). The inducer and noise segments were cross-faded over 20 ms using a raised cosine ramp. The noise segment replaced the real-world sound recording, so that the recorded sound and the noise were not physically present at the same time except during the onset and offset window. There were a total of 120 possible stimuli (24 sounds and 5 masker durations). To avoid priming effects, each participant heard each of the 24 sounds only once, in one of the conditions (see below).

To ensure that listeners understood the task, we also included practice and control trials where the inducer was higher in level than the noise and was either physically present or absent during the noise segment. We chose twenty-four additional sounds spanning the 4 categories described above. The same interrupting noise durations were also used. Stimulus construction was the same except that on the trials where the inducer was physically present, the noise was added to the inducer sound rather than replacing it. The level of the inducer relative to the noise was +6dB.

#### Procedure

Participants adjusted the playback level of their headphones to a comfortable setting using a calibration Gaussian noise stimulus. The level of the calibration noise was set to the maximum noise level that would appear in the main experiment.

The participants first performed a headphone check experiment [Woods et al., 2017]. If they passed the headphone check they then completed 18 practice trails with feedback. On each trial, participants reported whether they “heard the sound” or “didn’t hear the sound” during the noise. There were a total of 240 possible practice stimuli (24 sounds, 5 noise durations, and with the sound physically present or absent during the noise). The 18 practice trials were randomly selected from this set with the constraint that each sound was only presented once, noise durations were presented either 3 or 4 times, and that there were an equal number of trials where the sound was physically present or absent during the noise.

If performance on the practice trials exceeded 75%, participants proceeded to the main experiment. The task was the same as on the practice trials: participants again heard a 6s stimulus and reported whether they “heard the sound” or “didn’t hear the sound” during the noise. The main experiment consisted of 30 trials: 24 trials of the main experimental conditions and 6 control trials. The main experiment trials were a random subset of the 120 possible trials (24 sounds x 5 masker durations), with the constraint that each participant heard each of the 24 sounds once and each noise duration was presented 4 or 5 times. The sounds on the 6 control trials were distinct from the sounds in the main experimental trials.

#### Participants

160 participants attempted the experiment. 80 of these passed both the headphone check and the practice portion of the experiment (32 female, mean age = 37.4, s.d. = 12.2). Participants were unique to Experiment 2.

### Experiment 3 – Estimation of the extent of illusory continuity

#### Stimuli

We used twenty real-world sound recordings as the inducer sounds (Table 3). The stimuli were selected from the larger set used in Experiment 1 to span a range of sound stationarity and source types (e.g. textures, environmental sounds, speech, music and periodic sounds). Stimuli were 6 s in duration and began with a 2 s inducer followed by 4 s of white Gaussian masking noise. The 2 s inducer excerpt was randomly selected from a 7 s recorded sound. The level of the noise was set by the authors to plausibly mask the inducer (Table 3).

We included twenty control trials to ensure that participants could perform the task as instructed. The control stimuli consisted of an inducer that was higher in level than the noise and that ended either at the onset of the noise, or 1, 2, 3 or 4 s after the noise onset, continuing into the noise until this point (5 control conditions in total, Figure 4b). The inducer level was 55 dB SPL and was +6dB above the noise.

#### Procedure

Participants estimated the temporal extent by which the inducer perceptually continued into the noise. Following the 6 s stimulus, participants adjusted a slider to indicate the point in time where they heard the inducer end. During the stimulus presentation, participants were shown a progress bar plotting the temporal progression of the stimulus relative to the slider scale. The slider could be positioned from the noise onset (2-s) to the offset (6-s).

Participants completed 100 trials: 4 presentations of the 20 main experiment conditions along with 20 control trials. The pairings of control condition and inducer sound were randomized for each participant, with the constraint that each sound was presented once as a control. The order of trials was randomized for each participant.

Prior to the experiment, participants completed 20 practice trials that had the same structure as the control trials but with different condition/sound pairings. Feedback was provided to the participants after each trial, consisting of a visual indicator on the slider as to when the inducer sound ended.

#### Data Exclusion

Participants were included in the analysis if they reported the temporal extent of the inducer to within 250 ms of its actual extent on at least 85% of the practice trials. All ten participants satisfied this criterion and were thus included in the analysis (4 female, mean age = 22.4, s.d. = 2.1). These participants also completed the in-lab version of Experiment 1.

### Experiment 4a – Texture masking

The experiment was intended to reveal the level of noise required to mask a sound texture, for comparison to the effect of SNR on perceived continuity in Experiment 4b.

#### Stimuli

Twenty sound textures (Supplemental Table 4) were used in the experiment and were selected from a set of sounds used in previous studies of sound texture perception [McDermott et al. (2013)]. Each trial presented a 2 s sample of Gaussian noise and a superposition of a 2 s sample of Gaussian noise and a 2 s sound texture excerpt, in random order (Figure 5a – upper panel). We ran two versions of the experiment: one where the textures were excerpted from real-world recordings, and one where they were excerpted from synthetic textures. For the latter, the stimuli were random excerpts sampled without replacement from twelve 7 s samples synthesized from the statistics of each real-world texture (such that each trial was drawn from a unique exemplar; see Procedure).

The presentation level of the sound texture was fixed at 55dB SPL and the noise level varied from trial to trial, with an SNR range of −18dB and +12dB (in 6dB increments). Each stimulus was 2 s in duration, with an inter-stimulus-interval of 400 ms.

#### Procedure

Participants completed twenty blocks of trials, with one block for each of the twenty sound textures used in the experiment. Each block contained 2 presentations per SNR value, for a total of 12 trials, presented in random order. The experiment consisted of 240 trials in total. The order of the blocks was randomized for each participant.

At the beginning of a block participants were played a 2 s synthetic exemplar of the target texture without noise, so that they knew what to listen for. On each trial, participants heard the interval composed of noise plus the synthetic texture and the interval composed of only noise, in random order. Participants judged whether the target texture was present in the first or second interval. Every stimulus interval was generated from a unique noise segment and synthetic texture exemplar, such that the participants never heard the same waveform twice within or across trials. Participants had the option of refreshing their memory of the target texture by listening to it again in isolation at any point in the block.

#### Participants

Ten participants completed the experiment with real texture sounds (7 female, mean age = 20.7, s.d. = 1.8). A different group of ten participants completed the experiment with synthetic texture sounds (8 female, mean age = 22.4, s.d. = 2.71).

### Experiment 4b – Effect of masker level on texture continuity

#### Stimuli

The same twenty sound textures used in Experiment 4a were used in Experiment 4b. The stimuli consisted of sound textures interrupted with a noise segment (Figure 5a – lower panel). The stimulus was constructed by concatenating a 2-s excerpt of sound texture, followed by a 2 s Gaussian masking noise, followed by an additional 1s of the synthetic texture. The texture excerpts were the first 2 s and the last 1 s of a 5 s excerpt. Stimuli were constructed as in Experiment 1. We ran two versions of the experiment: one where the texture excerpts were taken from real-world recordings, and one where they were synthetic textures. For the latter, the stimuli were random excerpts sampled without replacement from twelve 7s samples synthesized from the statistics of each real-world texture. The presentation level of the sound texture excerpts was fixed at 55dB SPL. The SNR of the interrupting noise segment varied between −18dB and +12dB (in 6dB increments).

The experiment also included twenty control trials, one for each of the twenty sound textures, in which the texture was physically continuous during the interrupting noise segment, with an SNR value of +6dB.

#### Procedure

On each trial, participants heard a 5 s stimulus as described above. Participants judged whether the texture was continuous or discontinuous during the noise. The experiment consisted of 280 trials: 2 presentations of the 120 main experiment trials (6 masker levels x 20 textures) and 2 presentations of the 20 control trials. The order of the trials was randomized for each participant with the constraint that the same texture never occurred in consecutive trials.

#### Participants

The same ten participants from Experiment 4a completed Experiment 4b.

### Experiment 5 – Effect of gaps on texture continuity

This experiment measured the effect of temporal gaps on perceived texture continuity.

#### Stimuli

The same twenty textures from Experiment 4a/b were used in Experiment 5. The stimuli were constructed by interrupting a 5 s synthetic texture with 2 s of Gaussian noise or a combination of Gaussian noise and silence, the latter case resulting in a gap between the texture and the noise. The synthetic textures were random excerpts from one of eight 7-s samples synthesized from the statistics of each real-world texture. A different synthetic exemplar was used on each trial. The experiment had four conditions distinguished by the presence or absence of gaps: (i) no gaps (the sound texture and noise were contiguous), (ii) a gap before and after the noise (iii) a gap after the noise (iv) a gap before the noise (Figure 3C). The gaps were always 200 ms in length and replaced the beginning or end of the noise. The presentation level of the sound texture excerpt was set to 55dB SPL. The level of the intermediate noise segment was set individually for each sound as that which would mask the inducer texture were they presented simultaneously (validated by Experiment 4a). This level varied somewhat from texture to texture, but was always between 67.5 and 75 dB (corresponding to an SNR between −12.5 and −20dB; see Table 4).

In addition to trials with the main experimental conditions, we included 48 control trials in which the texture was physically present during the noise, to confirm that participants could perform the task. The control trials were constructed to mimic the response contours by modulating the amplitude of the synthetic texture during the intermediate noise segment (see schematics in Figure 6d). The stimulus consisted of a 2 s inducer texture, followed by a 2 s segment of “contoured” texture plus noise, followed by 1 s of texture. For the time-varying contour shapes, the contoured texture was constructed by multiplying the texture by a raised cosine envelope that varied between 0 and 1 with a phase set to yield the desired shape. The SNR value between the unattenuated inducer texture and noise was +6dB, such that the texture was audible when superimposed on the noise (compare to Figure 2b, left panel). The portion of the synthetic texture corresponding the response contour extended for 2 s and co-occurred with the noise segment. The six contour shapes were presented twice for each of the four experiment conditions, with the texture randomly selected from the 20 textures used in the main experiment. For the control conditions with a silent gap, the first and/or last 200 ms of the texture+noise combination was replaced with silence, such that each contour was presented in each of the four stimulus configurations, to better assess whether participants could identify the contour shape across the various gap arrangements.

#### Procedure

On each trial participants heard a 5 s stimulus and selected one of 6 response contours to indicate what they heard: continuous throughout, fade-out, present at beginning and end of the noise (‘dip’), present in the middle of the noise (‘glimpse’), fade-in, and absent throughout. Prior to the main experiment, participants performed 24 practice trials, which had the same form as the control trials, but with combinations of textures and conditions that were not used in the control trials. Participants received feedback for the practice trials.

The experiment consisted of 232 trials: two presentations of the 4 main conditions with each of the 20 textures, 48 control trials, and the initial 24 practice trials. The order of the practice trials was randomized but always occurred at the beginning of the experiment. The main experiment trials and control trials were randomized.

#### Participants

The same ten participants from Experiment 4a and 4b (version with synthetic sound textures) completed Experiment 5.

### Experiment 6 – Statistical integration of illusory texture

#### Texture Step and Morph Synthesis

Synthetic texture stimuli were generated using a variant of the McDermott and Simoncelli^14^ sound texture synthesis system. As introduced in a recent publication^20^, the original texture synthesis system was modified to facilitate the generation of “texture morphs” (sound textures generated from statistics sampled at points along a line between two textures) and “texture steps” (sound textures that underwent a change in their statistics at some point during their duration).

We measured the statistics of 50 real-world texture recordings, averaged their statistics, and used these average statistics to define the “mean” texture. The statistics measured from individual real-world texture recordings defined a “reference” texture that varied across blocks. The cochlear envelope means (*μ*_*k*_) of the reference were set equal to those for the mean texture, such that spectral content was constant across stimuli (so that any integration effects would be likely to reflect higher order statistics). We synthesized texture morphs by sampling discrete points along a line in the space of statistics between the mean and the reference. We generated texture steps so that their statistical properties changed at some point in time by stepping from one set of texture statistics to another. For the texture steps, we first synthesized a texture from one set of statistics. We then sampled a second set of statistics from the line between the mean and reference, using the first synthetic texture as the seed to the synthesis system. The resulting synthetic textures were then windowed (with rectangular windows) and summed to yield a texture step with the desired change in statistics that occurred at a specific point in time. This procedure produced signals whose statistics changed over time without introducing artifactual discontinuities that might otherwise result from concatenating signals with different statistics.

#### Stimuli

Four reference textures were used (River running over shallows, Pneumatic drill, Applause, and Lawn edger). These textures were selected because they elicited a consistently high continuity response when interrupted with masking noise (refer to Experiment 4b). Each trial presented a “step” stimulus followed by a “morph” stimulus.

The step stimulus statistics began at either 25% or 75% of the distance between the mean and reference, and stepped to the 50% point between the mean and reference. The step stimulus was 5 s in duration. There were conditions, each with a different type of step stimulus, each of which had two step directions (beginning at either the 25% or 75% point; Figure 4C). The first two conditions included a step positioned either 1 s or 3 s from the endpoint. The third condition replaced an intermediate segment of the 3 s step with 2 s of Gaussian masking noise, ending 1 s from the endpoint. The fourth condition was identical to the third conditions, except that the first 200 ms of the masking noise was replaced by silence. In all conditions, the texture was not physically present during the interrupting noise. Five exemplars were synthesized for each step condition for each reference texture.

The second stimulus on a trial (the morph) was generated with statistics drawn from one of 10 positions between the mean and the reference (0, 0.25, 0.35, 0.4, 0.45 0.55, 0.6, 0.65, 0.75, and 1, where the number denotes the fractional position between mean and reference, with 0 denoting the mean and 1 denoting the reference). The morph duration was two seconds. The step and morph were separated by an inter-stimulus-interval of 250 ms. The sound texture portion of the signal was presented at 55dB SPL. When present, the masking noise was set between 14-18 dB above the level of the texture (Table 2), with the exact level set individually for each texture as that necessary to mask the texture when the noise and texture were superimposed.

#### Procedure

Trials were blocked by the reference texture. Each block presented one trial for each of the 10 morph positions paired with a randomly selected step condition. Each step condition occurred once with each morph position for each reference texture across the experiment. The order of the blocks was always random subject to the constraint that two blocks with the same reference texture never occurred consecutively. The stimulus used on a trial was randomly selected from five pre-generated synthetic textures for each condition and reference texture. At the start of each block participants were played a 5s excerpt of the reference texture, which they could listen to as many times as they wished. Participants had the option of listening to the reference texture again between trials during the block.

Participants selected the interval that was most similar to the reference texture. Participants were informed that the step interval could change over time and could contain an interrupting noise segment. They were instructed to base their judgments on the endpoint of the step interval, a task that we validated in a previous publication^20^. Feedback was not provided.

Participants completed a practice session consisting of 4 blocks (each with a different randomly selected reference texture) of 6 trials to familiarize participants with the task and the reference sounds. In the practice session the first stimulus did not contain a step, and was instead generated with constant statistics drawn from the midpoint between the reference and mean, and the second stimulus was generated from statistics drawn from one of six positions (0 - mean, 0.25, 0.4, 0.6, 0.75, 1 - reference). Both stimuli were 2-s in duration. Feedback was given following each practice trial.

#### Participants

To restrict the analysis to participants who could perform the task (which was more difficult than the others used in this paper), we excluded participants who did not perform at least 85% correct when the morph was set to the statistics of the reference or mean (i.e., the most discriminable difference used in the experiment) for at least one of the step conditions. Sixteen participants completed the experiment, of which thirteen met the inclusion criterion (9 female, mean age = 21.2, s.d. = 1.5). The participants were unique to this experiment, and did not participate in any other experiments.

### Experiment 7 – Illusory continuity of sounds in auditory scenes

#### Stimuli

We selected 20 sound textures and 20 non-textures from a larger set as those that had high and low values of sound stationarity (Figure 8a). Each stationary sound (texture) was paired with a variable sound (non-texture). The authors attempted to pair the sounds such that they could plausibly occur in natural listening environments (e.g. applause paired with music; see Table 4).

Each trial began with a 2 s cue sound, followed by a 400 ms silent gap, followed by a 5 s stimulus. There were three conditions, differentiated by the cue sound: (i) texture, (ii) non-texture, and (iii) superposition of texture and non-texture. The 5 s stimulus was the concatenation of 2 s of the superposition of texture and non-texture (the inducer), followed by 2 s of Gaussian noise (the masker), followed by another 1 s of the superposition of texture and non-texture. The sound segments were cross-faded over 20 ms with a raised cosine ramp. The waveform of the cue and initial 2 s of the texture, non-texture, or their superposition were identical. The noise level was set by the authors to plausibly mask the texture + non-texture inducer (Table 6).

The experiment also included forty control trials to confirm task comprehension and compliance. Similar to the main experimental trials, the control trials cued participants to report the continuity of a sound: either the texture, the non-texture or the combined sound. But unlike the main experimental trials, the inducer in the control trials was higher in level than the noise and the cued sound was either physically present (continuous) or absent during the noise (Supplemental Figure 6). The texture and non-texture in the inducer were equal in level, and the inducer (texture + non-texture) was 55 dB SPL. The noise was set to 6 dB below the cued sound (either the texture, non-texture or their combination, depending on the condition). The cue was the same level as the corresponding source signal(s) within the inducer. There were 5 continuous control conditions, where the cued sound was physically present during the noise, but where the un-cued sound was present or absent during the noise depending on the condition. There were also 5 discontinuous control conditions, where the cued sound was physically absent during the noise, but where the un-cued sound was present or absent during the noise depending on the condition. If listeners were reporting the subjective presence of the cued sound (as instructed), they should report continuity on the 5 continuous control conditions but not the 5 discontinuous control conditions. The results of the 5 continuous control conditions and the 5 discontinuous control conditions were analyzed together (Supplemental Figure 6). There were a total of 200 possible control trials (10 conditions crossed with 20 inducer sound pairs).

#### Procedure

On each trial, participants heard the cue sound followed by the stimulus. Participants judged whether the cued sound was continuous or discontinuous during the interrupting noise. Participants were instructed to respond with continuous only if the cued sound continued during the noise. For instance, if the cue sound was the sound pair, the listeners would respond with “continuous” if the sound pair continued during the noise and “discontinuous” of only one sound continued or neither sound continued during the noise. Participants performed 160 trials in total: 120 trials of the three main conditions and 40 control trials. Each of the three main conditions occurred twice for all twenty sounds. The control trials were randomly selected for each participant from the set of 200 possible stimulus/condition pairings, with the constraint that each inducer sound pair occurred twice and each condition occurred four times. The order of the trials was randomized for each participant.

Prior to the main experiment, participants performed 20 practice trials with feedback. The practice trials were similar in structure to the control trials, but with distinct stimulus/condition pairings. The feedback indicated whether the cued sound was in fact continuous or discontinuous during the noise.

#### Data Exclusion

Participants were included in the analysis if they accurately reported the presence of the cued inducer during the noise in at least 85% of the practice trials. All ten participants met this criterion and were included in the analysis (5 female, mean age = 23.1, s.d. = 1.9). These participants were exclusive to Experiment 7.

### Sample Sizes

The sample size for each experiment was chosen to yield stable results based on split-half reliability of the results in pilot experiments.

#### Experiment 1 (Illusory continuity for a large set of natural sounds)

##### In-lab

We ran a pilot version of the experiment (N=8). The sound set was slightly different from that of Experiment 1. We calculated the split-half reliability (Pearson’s correlation coefficient) of the main result (proportion of continuous responses for each of the inducer sounds) for sample sizes ranging from N = 2 to 8 with random splits of the participants (N=10,000). We found the split-half reliability increased with sample size from 0.60 (N = 2) to 0.87 (N = 8). We extrapolated that a sample size of 13 would have a likely test-retest correlation greater than 0.9, and chose this as our sample size.

##### Online

We used the power analysis for Experiment 2, which used a similar task and was also conducted online. This yielded a sample size of 80, which we slightly exceeded.

#### Experiment 2 (Illusory continuity across different masker durations)

We ran a pilot version of the experiment (N=44). The experiment differed from Experiment 2 in two respects: (i) participants completed all sound/condition pairings, resulting in multiple trials with the same sounds, and (ii) the sound set was slightly different. We performed a power analysis by selecting subsets of 24 trials (to match the number of trials in Experiment 2). We measured the Pearson’s correlation coefficient between random splits of participants for sample sizes ranging from N = 2 to 44. The split-half reliability increased with sample size from 0.17 (N = 2) to 0.84 (N = 44). We extrapolated that a sample size of 80 would give an expected split-half reliability greater than 0.9, and chose this as the sample size.

#### Experiment 3 (Estimation of the extent of illusory continuity)

We ran a pilot experiment (N=6). The pilot experiment included 40 sounds (rather than the 20 sounds used in Experiment 3), and the noise extended for only 2s after the inducer (rather than the 4s used in Experiment 3). Split-half reliability increased with sample size from 0.47 (N = 2) to 0.83 (N = 6). We chose a sample size of 10 to yield an expected reliability in excess of 0.9.

#### Experiment 4a (Texture masking)

We ran a pilot version of the experiment (N = 9) with real-world sound textures that used a slightly different set of textures and a reduced range of SNR conditions (spanning −18dB to +6dB in 6dB increments). We calculated the split-half reliability (Pearson’s correlation coefficient) of the main result (proportion correct for each of the SNR conditions) for sample sizes ranging from N = 2 to 8 with random splits of the participants (N=10,000). The mean split-half reliability increased with sample size from 0.86 (N = 2) to 0.96 (N = 8). We selected a target sample size of 10 as that which would likely yield a split-half correlation of at least 0.9.

#### Experiment 4b (Effect of masker level on texture continuity)

We ran an analogous pilot version of Experiment 4b. Using the same power analysis procedure described for Experiment 4a, we found the split-half reliability increased with sample size from 0.71 (N = 2) to 0.91 (N = 8). We selected a target sample size of 10 as that which would provide an expected reliability correlation greater than 0.9.

#### Experiment 5 (Effect of masker contiguity on texture continuity)

We ran a pilot version of the experiment (N = 8). The experiment was identical to Experiment 5 except that the inducer sounds were real-world texture recordings and the sound set was slightly different. We calculated the split-half reliability (Pearson’s correlation coefficient) of the mean results (the proportion responses for each contour shape) for sample sizes ranging from N = 2 to 8 with random splits of the participants (N=10,000). The split-half reliability increased with sample size from 0.73 (N = 2) to 0.91 (N = 8). We selected a sample size of 10 as that which would provide an expected test-retest correlation greater than 0.9.

#### Experiment 6 (Statistical integration of illusory texture)

We had performed power analyses for this type of experiment in a previous publication^20^. We chose a sample size of 10 as this yielded an expected split-half reliability of the mean results that exceeded 0.9.

#### Experiment 7 (Illusory continuity of sounds in auditory scenes)

The methodology of this experiment was similar to that of the in-lab version of Experiment 1, and we targeted the same sample size identified for that experiment (N=10) for an expected reliability in excess of 0.9.

### Statistics and Data Analysis

Data was assumed to be normally distributed and was evaluated as such by eye. All reported correlations are Pearson’s correlations.

#### Experiment 1 (Illusory continuity for a large set of natural sounds)

We evaluated the reliability of the online data by computing Pearson’s correlation coefficient between the mean proportion of continuous responses for splits of participants (using 10,000 random splits). The reported correlation is the mean over the resulting 10,000 correlation values.

#### Experiment 2 (Illusory continuity across different masker durations)

The effects of sound category and noise duration were assessed using a multi-level repeated-measures ANOVA.

#### Experiment 3 (Estimation of the extent of illusory continuity)

A repeated-measures ANOVA was used to evaluate the variation in the temporal extent of continuity across inducer sounds.

#### Experiment 4a/b (Texture masking and continuity)

Repeated measures analyses of variance (ANOVA) were used to test for effects of SNR on masking and continuity judgments.

To compare the threshold SNRs that produced masking and illusory continuity (as shown in Supplemental Figure 2), we fit logistic functions to the data from each experiment. Using these fits, we defined the masking threshold as the SNR producing a proportion correct of 0.833 in the texture masking experiment (4a) and the continuity threshold as the SNR producing judgments of continuity on 0.667 of the trials for texture continuity experiment (4b). These values represent 2/3 of the dynamic range of the psychometric functions (the masking curve spans 0.5 and 1, whereas the continuity curve spans 0 and 1). We compared the threshold SNR values for masking and illusory continuity with a Spearman’s rank correlation coefficient across sounds (Supplemental Figure 2b)

#### Experiment 5 (Effect of gaps on texture continuity)

Single sample or paired two-tailed *t*-tests were used to compare results for individual conditions to chance, and for pairwise comparisons between conditions, respectively.

#### Experiment 6 (Statistical integration of illusory texture)

We fit psychometric (logistic) functions to the mean results for each step condition and step direction (i.e. the curve plotting the proportion of trials where the morph was judged to be more similar to the reference, for each morph position), and from the fitted functions obtained the point of subjective equality for each step condition and direction. We quantified the step-induced bias as the difference between the points of subjective equality for the two step directions for each condition. Confidence intervals on these bias measurements were derived by bootstrap (10,000 samples)^20^. Statistical significance of differences in the bias between conditions was estimated from the bootstrap distributions of the bias by fitting a Gaussian and then computing the p-value from the Gaussian.

#### Experiment 7 (Illusory continuity of sounds in auditory scenes)

Differences between conditions were evaluated with paired two-tailed *t-tests*. An un-paired, two-tailed *t-test* was used to evaluate differences in continuity for the same two sound categories (textures and non-textures) between Experiment 7 and Experiment 1.

## Supplemental Material

**Table 1.** Sounds used in Experiment 1. Masking noise SNR (dB) indicates the relative difference in amplitude between the inducer sound and the Gaussian masking noise, selected by the first author to produce masking of the texture by the noise when they were superimposed.

| Inducer | Masking noise SNR (dB) | Inducer | Masking noise SNR (dB) |
| --- | --- | --- | --- |
| Bees | -18 * | Pneumatic drill | -14 |
| Birds in spring | -16.5 | Pneumatic rills (road-works) | -14 |
| Cat lapping milk | -14.5 | Pouring coffee beans | -17.5 |
| Frogs | -12.5 | Pouring coins | -16 |
| Swamp insects | -16 | Static | -16 |
| Rattlesnake | -16.5 | Scratching beard | -17 |
| Sparrows (large excited group) | -16.5 | Shaking paper | -15 |
| Summer day in the south | -12.5 | Ship anchor up | -13.5 |
| Swamp noises | -16.5 | Snare steel sequence | -23 |
| Swarm of bees | -17.5 | Radio static | -12.5 |
| Fire | -18 | Steam train slow | -20 |
| Heavy rain on hard surface | -17 | Tambourine shaking | -14 |
| Wind whistling | -17.5 | Tapping rhythm | -16 |
| Rain beating against window | -13.5 | Train | -19 |
| Rain | -14.5 | Windshield wipers | -20 |
| AC unit | -23 | Animated crowd in a large hall | -13.5 |
| AM radio noise | -14 | Rhythmic applause | -19 |
| Blender | -16.5 | Applause in large hall | -18 |
| Brushing teeth | -15 | Applause | -15 |
| Car interior | -17.5 | Babble | -18 |
| Church bells | -15.5 | Enthusiastic applause | -17.5 |
| City room teletype | -16 | German rail station announcer | -21.5 |
| Coin rolling on plate | -14.5 | Large railway station | -15.5 |
| Crumpling paper | -17.5 | Men marching | -17 |
| Crunching cellophane | -14 | Music (apache) | -21.5 |
| Dinner triangle | -17.5 | Music (bluegrass) | -20.5 |
| Drumroll long | -19 | Music (orchestra) | -21.5 |
| Lawn edger | -15.5 | Speech (German female) | -23.5 |
| Electric drill | -16.5 | Speech (German male) | -23 |
| Firecrackers | -17 | Bathroom sink | -15.5 |
| Frying bacon | -14.5 | Bubbles | -23 |
| Galloping horse | -24 | Bubbling water | -19 |
| Hammering copper | -18 | Cave drop | -20.5 |
| Hand sanding | -12.5 | Fast running river | -20 |
| Helicopter | -21.5 | Ocean wave slow | -21.5 |
| Jingling coins | -16 | River running over shallows | -18 |
| Jogging on gravel | -12.5 | Stream small | -19 |
| Manual typewriter | -14 | Water dribbling | -20 |
| Modulated radio static | -14.5 | Water lapping gently | -19.5 |
| Newspaper printing press | -11.5 | Water movement | -21 |

\* Negative SNR value corresponds to the noise (masker) being higher in level than the signal (inducer).

**Table 2.**
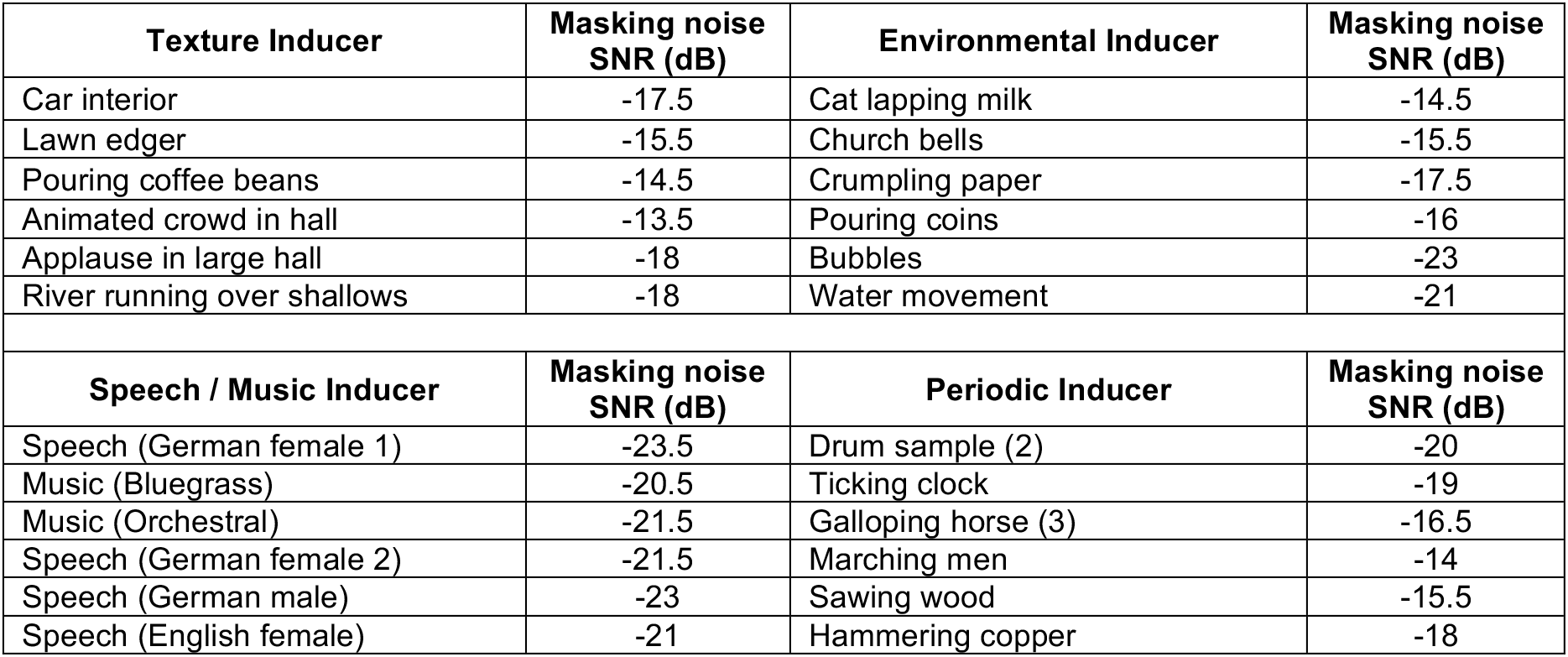
Categories and sounds used in Experiment 2 (varying noise duration).

**Table 3.** Sounds used in Experiment 3 (extent estimation task).

| **Inducer** | **Masking noise SNR (dB)** | **Inducer** | **Masking noise SNR (dB)** |
| --- | --- | --- | --- |
| Swamp insects | -16 | Rain beating against window | -13.5 |
| Car interior | -17.5 | Firecrackers | -17 |
| Jogging on gravel | -12.5 | Shaking paper | -15 |
| Applause in large hall | -18 | Speech (German female 1) | -23.5 |
| Music (Bluegrass) | -20.5 | Bubbling water | -19 |
| Rattlesnake | -16.5 | Brushing teeth | -15 |
| Lawn edger | -15.5 | Helicopter | -21.5 |
| Pouring coffee beans | -17.5 | Tapping rhythm | -16 |
| Babble | -18 | Large railway station | -15.5 |
| Speech (German male) | -23 | Water movement | -21 |

**Table 4.** Sounds used in Experiments 4a, 4b, and 5 (masking vs. continuity, and effect of gaps). Masking Noise SNR was that used in Experiment 5. The numbers in parentheses are the masking thresholds measured in Experiment 4a.

| Inducer | Masking noise SNR [dB (measured)] | Inducer | Masking noise SNR [dB (measured)] |
| --- | --- | --- | --- |
| Rattlesnake | -16.5 (-6.5) | Pneumatic drill | -14 (-2.5) |
| Heavy rain on a hard surface | -17 (-4.5) | Pneumatic drills (road works) | -14 (-4.5) |
| Wind whistling | -17.5 (-10.5) | Pouring coffee beans | -17.5 (-16.5) |
| Rain | -14.5 (-6.5) | Ship anchor | -13.5 (-3.5) |
| Blender | -16.5 (-9.5) | Radio static | -12.5 (-4) |
| Car interior | -17.5 (-11.5) | Babble (large hall) | -18 (-8.5) |
| Crunching cellophane | -14 (-7) | Applause in large hall | -18 (-5.5) |
| Lawn edger | -15.5 (-9.5) | Applause | -15 (-6.5) |
| Electric drill | -16.5 (-8) | Fast running river | -20 (-3.5) |
| Frying bacon | -14.5 (-8.5) | River running over shallows | -18 (-7.5) |

**Table 5.** Sounds used in Experiment 6 (texture step experiment). Masking noise SNR was used for all stimuli for a given reference texture.

| Reference | Masking Noise SNR (dB) |
| --- | --- |
| River running over shallows | -18 |
| Pneumatic drill | -14 |
| Applause | -16 |
| Lawn edger | -17.5 |

**Table 6.**
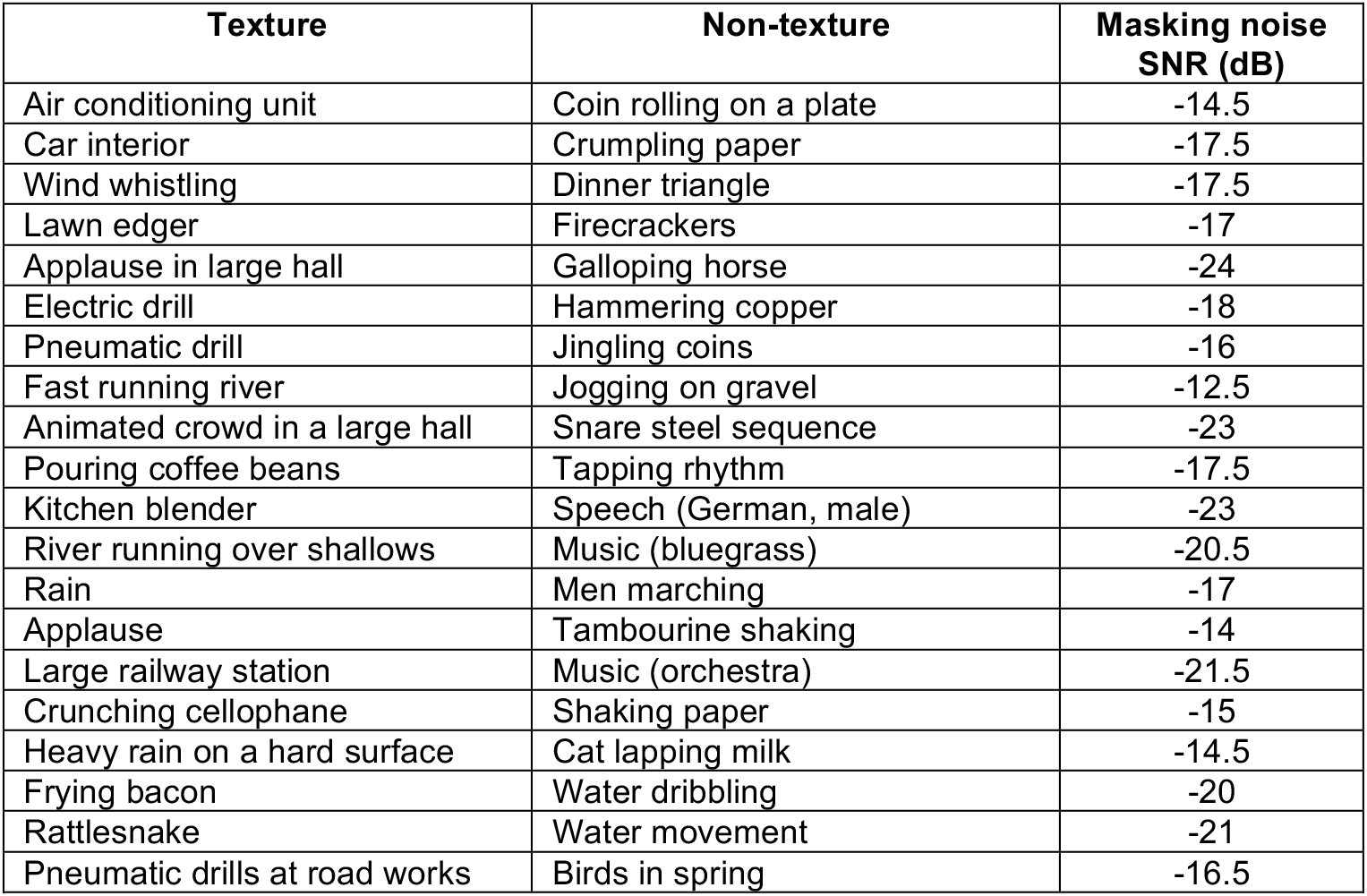
Sounds used in Experiment 7 (texture continuity with concurrent non-textures).

**Supplementary Figure 1.**
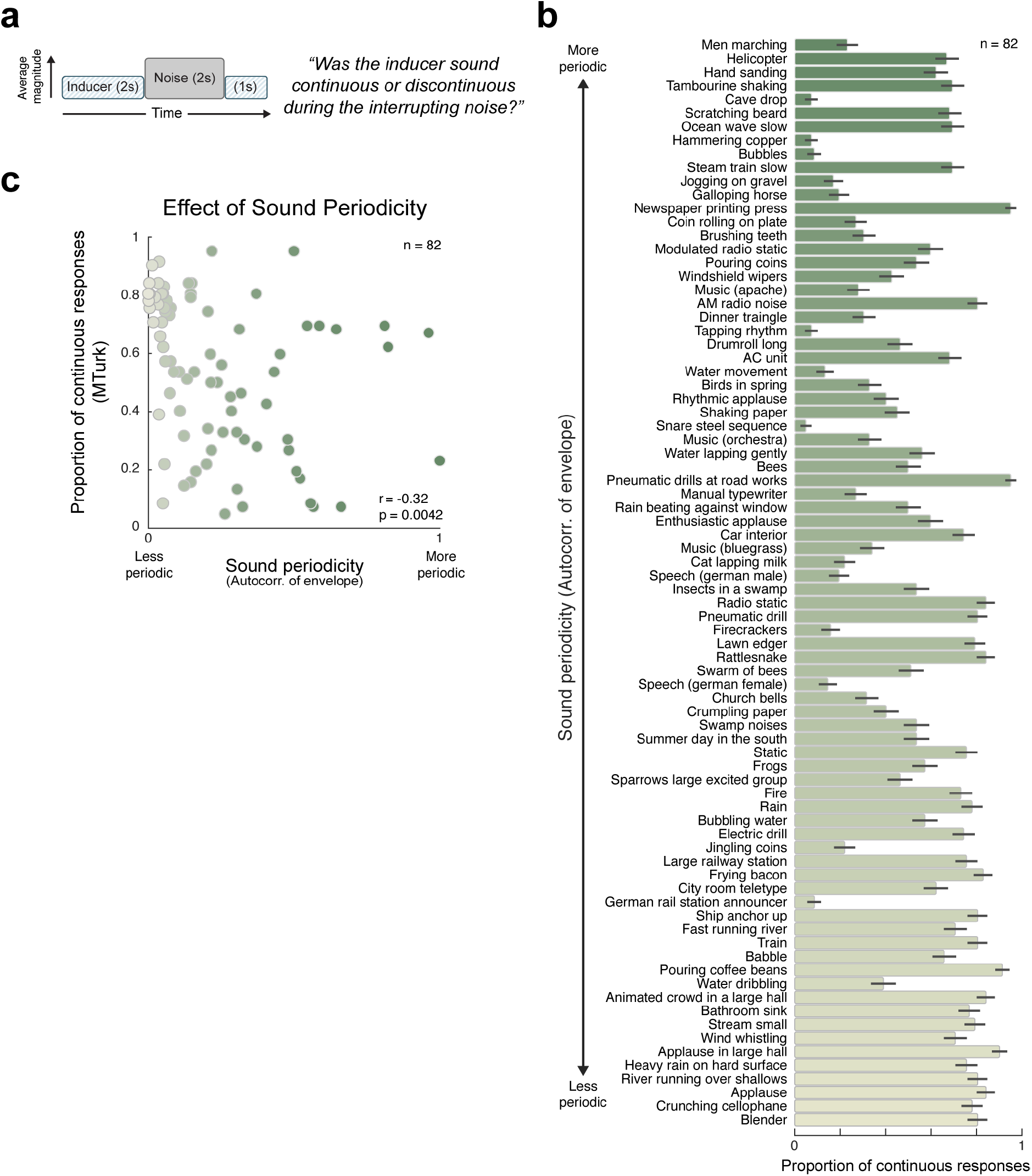
Results of Experiment 1 ordered by sound periodicity. **(a)** Schematic of stimulus condition of Experiment 1. **(b)** Results of Experiment 1. Like Figure 2f except that sounds are sorted by their periodicity, and color denotes periodicity. Periodicity was computed from the autocorrelation of the Hilbert envelope of the sound waveform, downsampled to 400 Hz. The periodicity measure was the height of the largest autocorrelation peak for lags between 0.125 s and 0.5 s, normalized by the autocorrelation at lag 0. Error bars show SEM. **(c)** Mean proportion of continuous responses plotted as a function of sound periodicity.

**Supplementary Figure 2.**
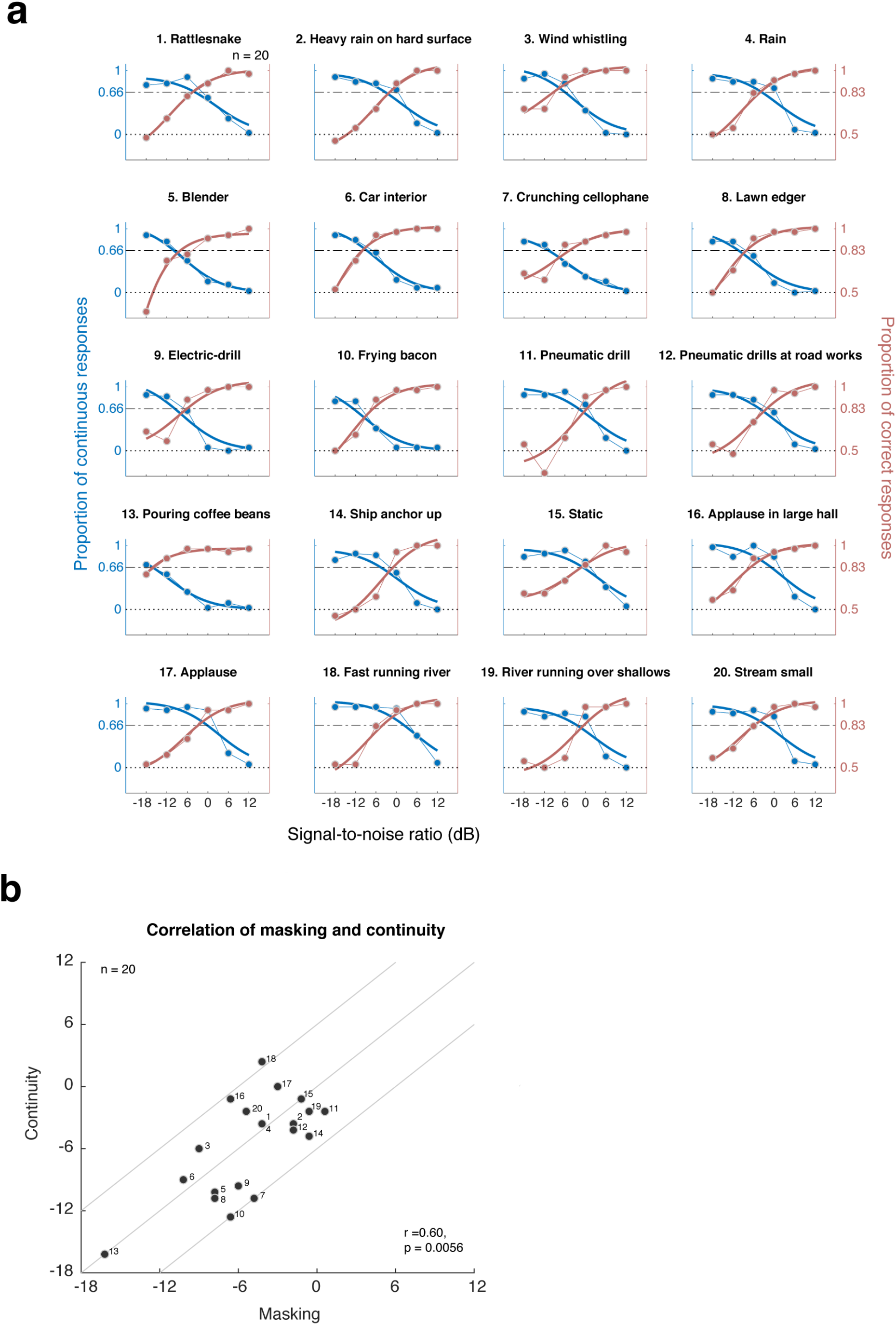
Results for the masking and continuity experiments (4a/4b) plotted separately for individual sounds. To increase power, this analysis combined the data from Experiments 4a/4b with that of a pilot experiment (n=10) that was identical except that the stimuli for a given reference sound were generated from a single synthetic texture exemplar (yielding n=20 in total). **(a)** Masking and continuity curves for individual sounds. The data points and lite/thin lines show the mean response across SNR values. The dark/thick lines show logistic function fits. The horizontal dashed line shows the threshold (masking 0.833, continuity 0.666) value used to relate masking and continuity in b. **(b)** Spearman correlation between masking and continuity thresholds. Index refers to sounds in subplot a. Diagonal lines show 0dB and +/− 6dB.

**Supplementary Figure 3.**
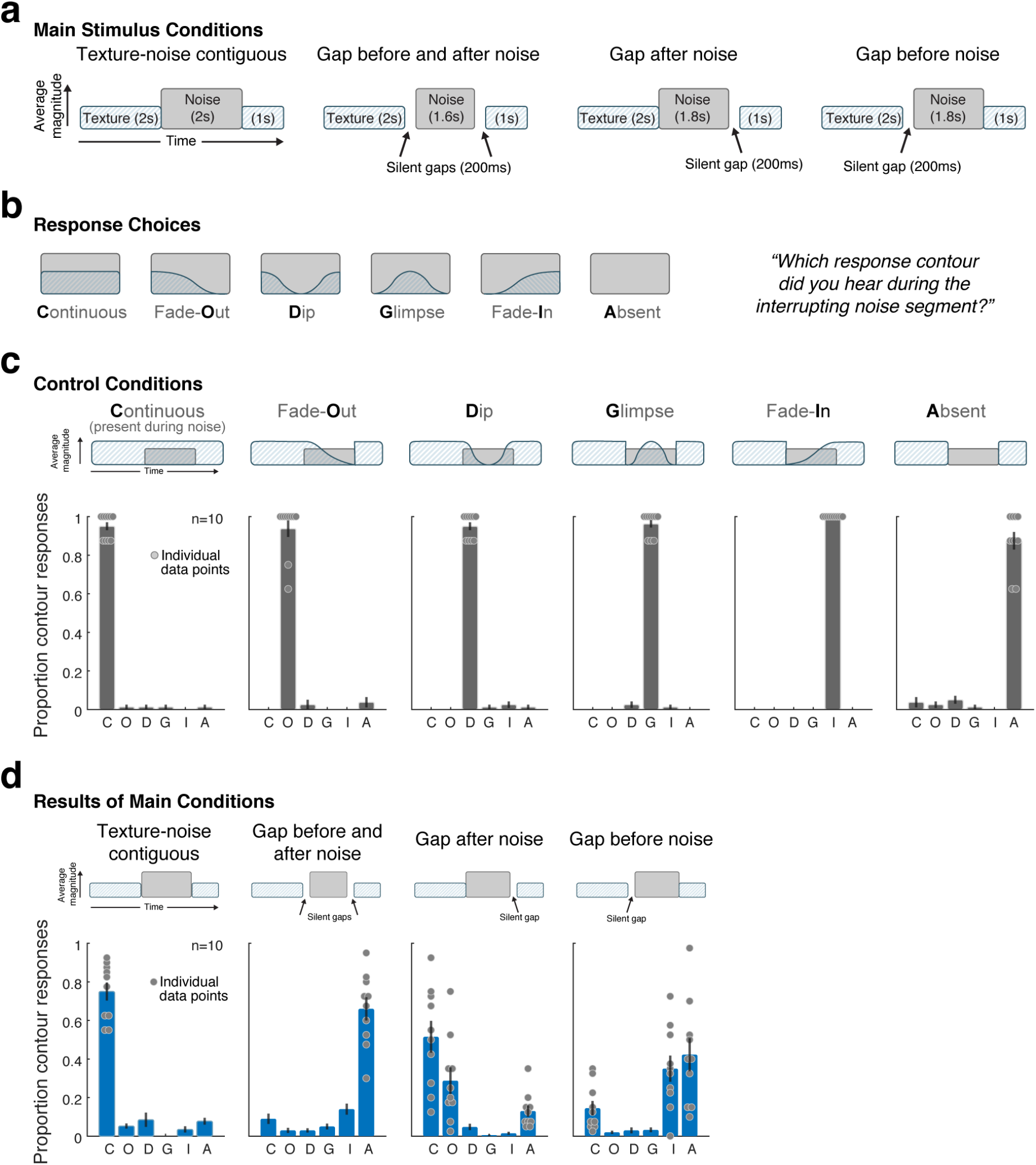
Replication of Experiment 5 with real-world sound texture recordings. (caption duplicated from Figure 6 caption) **(a)** Listeners heard a synthetic inducer texture interrupted with masking noise and reported their perceptual experience during the interrupting noise segment. The experiment included 4 conditions, differing in the contiguity of the texture and noise (via silent gaps inserted before and/or after the noise). See Supplementary Figure 3 for analogous experiment with real-world texture recordings (which yielded similar results). **(b)** Listeners chose one of six response contours to describe their perceptual experience during the interrupting noise segment. The contour response code is indicated as the bold letter for each response (e.g. “C” for “**C**ontinuous”). **(c)** To confirm task comprehension/compliance, the experiment included control trials where the texture was physically present during the intermediate noise segment and amplitude modulated according to one of the response contours. The stimulus for each condition is schematized above each of the six subplots. Graphs plot the proportion of trials on which each response was chosen. Here and elsewhere the error bars show SEM. Data for individual participants is plotted as dots for the response choices selected above chance levels for each condition. **(d)** Results of main experimental conditions. Each subplot corresponds to a condition (shown schematically above). Data for individual participants is plotted as dots for the response choices selected above chance levels for each condition.

**Supplementary Figure 4.**
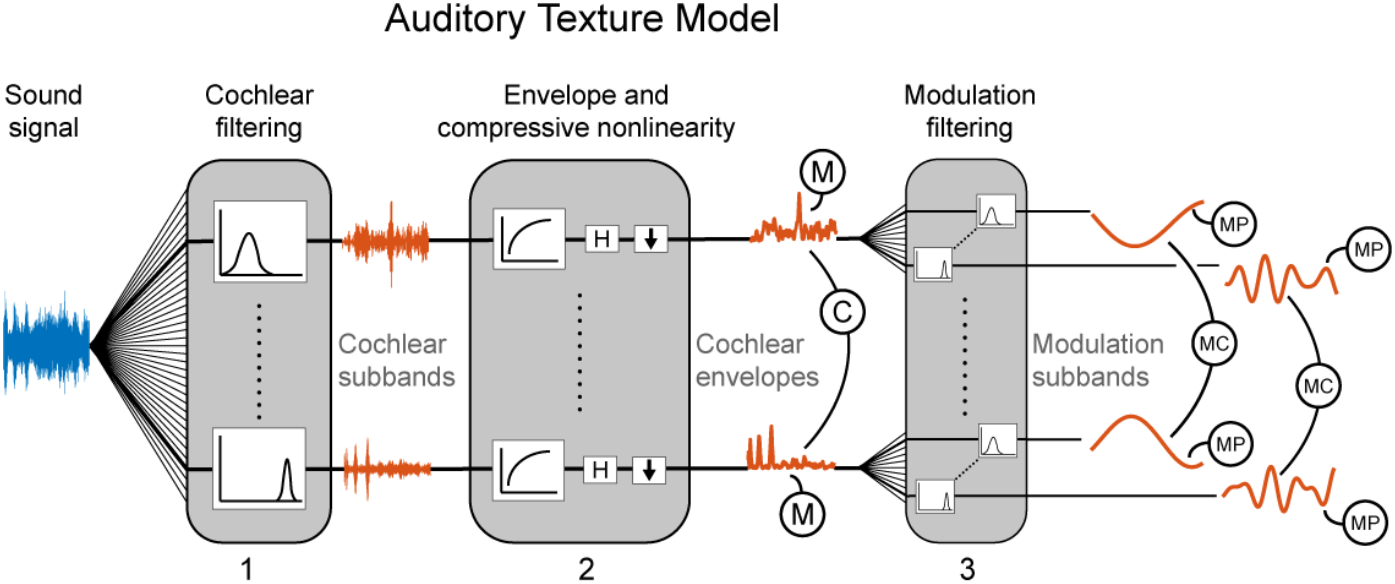
Auditory texture model. The model was adapted from that of McDermott and Simoncelli (2011). Statistics are measured from an auditory model capturing the tuning properties of three stages of the peripheral and subcortical auditory system. The cochlear envelope marginal statistics (M) comprise the mean, coefficient of variance, skewness and kurtosis. Pair-wise envelope (C) correlations were computed between neighboring cochlear envelope bands. The modulation subband statistics comprise the modulation power (MP; the variance of the modulation normalized by the corresponding total cochlear envelope variance) and modulation correlations (MC) between modulation subbands.

**Supplementary Figure 5.**
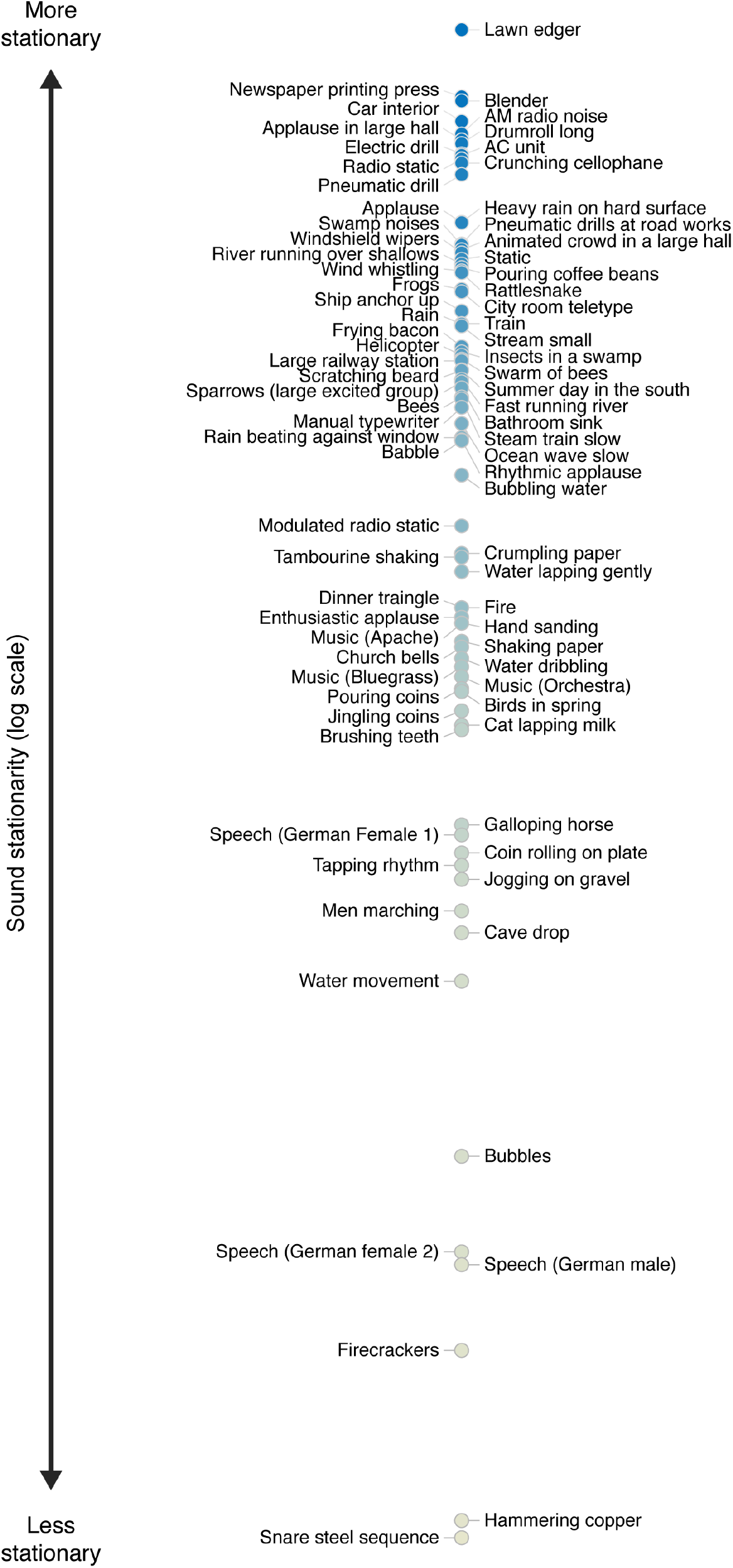
Stationarity of sounds used in Experiment 1 (80 sounds).

**Supplementary Figure 6.**
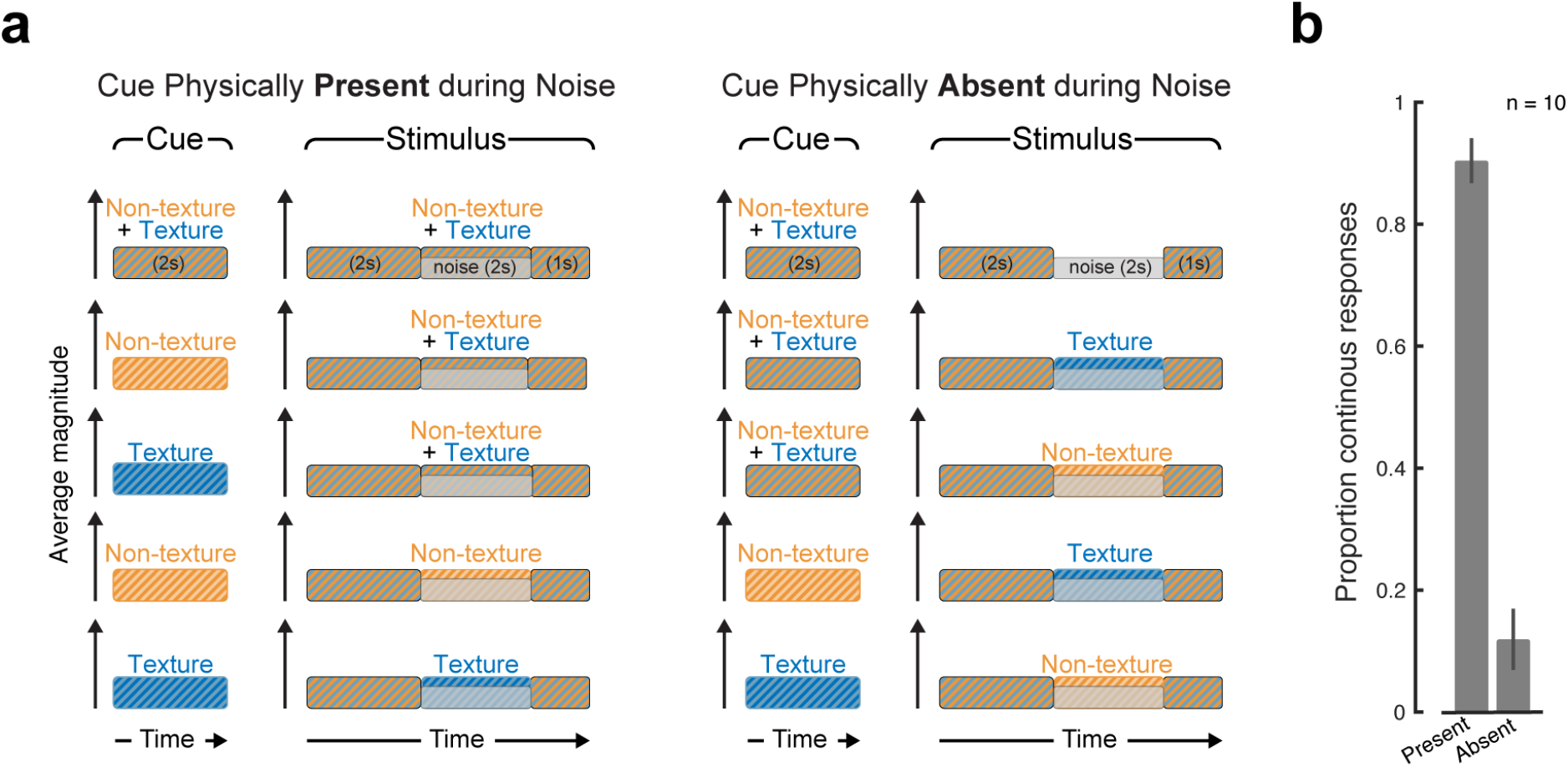
Control conditions for Experiment 7. (a) The stimuli varied in the sounds that were physically present during the noise. In five of the conditions the cued sound was physically present and in the other five the cued sound was physically absent during the noise segment. (b) Results for control conditions shown in a. Proportion of trials on which participants judged the cued stimulus to continue during the noise, averaged within the two groups of conditions (present or absent). Error bars show SEM. The results indicate that participants were performing the task as intended, in that they reported perceptual continuity when the cued sound was physically continuous, but not when it was unambiguously absent.

## Acknowledgements

The authors thank the McDermott laboratory for comments on the manuscript, in particular Maddie Cusimano and Jenelle Feather for comments on the penultimate draft. This work was supported by a McDonnell Foundation Scholar Award to J.H.M. and NIH Grant No. 1R01DC014739-01A1. The funding agencies were not otherwise involved in the research, and any opinions, findings and conclusions or recommendations expressed in this material are those of the authors and do not necessarily reflect the views of the McDonnell Foundation, or NIH.

## Author contributions

R.M. and J.H.M. designed the experiments. R.M. implemented experiments, and collected and analyzed the results. R.M. and J.H.M. wrote the paper.

